# Pangenome Analysis of Enterobacteria Reveals Richness of Secondary Metabolite Gene Clusters and their Associated Gene Sets

**DOI:** 10.1101/781328

**Authors:** Omkar S. Mohite, Colton J. Lloyd, Jonathan M. Monk, Tilmann Weber, Bernhard O. Palsson

## Abstract

The growing number of sequenced genomes enables the study of secondary metabolite biosynthetic gene clusters (BGC) in phyla beyond well-studied soil bacteria. We mined 2627 enterobacterial genomes to detect 8604 BGCs, including nonribosomal peptide synthetases, siderophores, polyketide-nonribosomal peptide hybrids, and 60 other BGC types, with an average of around 3.3 BGCs per genome. These BGCs represented 212 distinct BGC families, of which only 20 have associated products in the MIBiG standard database with functions such as siderophores, antibiotics, and genotoxins. Pangenome analysis identified genes associated with a specific BGC encoding for colon cancer-related colibactin. In one example, we associated genes involved in the type VI secretion system with the presence of a colibactin BGC in *Escherichia*. This richness of BGCs in enterobacteria opens up the possibility to discover novel secondary metabolites, their physiological roles and provides a guide to identify and understand PKS associated gene sets.

## Main

Secondary metabolites produced by a range of microorganisms display medicinally and industrially important properties, as well as mediate microbe-host and microbe-microbe interactions. Secondary metabolite biosynthesis often involves mega-enzymes such as polyketide synthases (PKS) and non-ribosomal peptide synthetases (NRPS) that are encoded by large biosynthetic gene clusters (BGCs). Recent advances in genome sequencing technology and genome mining tools revealed an unexplored richness and diversity of BGCs encoding secondary metabolites ^1–4^. In addition, the availability of a large number of genomes from the same species allowed for pangenome analysis revealing intra-species diversity, such as metabolic capabilities ^5,6^. The focus of many genome mining based studies have been well-established secondary metabolite producers, such as bacilli, actinobacteria, or myxobacteria ^7,8^. In comparison with many of the popular secondary metabolite producers, *Escherichia coli* or other enterobacteria have larger availability of sequenced genomes, higher quality of genome annotations, comprehensive curated databases, and extensive tools for data analysis. With the exception of *Photorhabdus*, *Xenorhabdus* and related genera, which are known to produce a diverse range of secondary metabolites ^9,10^, enterobacteria are known to produce few secondary metabolites. These include metal ion chelators like enterobactin, yersiniabactin ^11,12^, colon cancer-related genotoxin colibactin ^13,14^, antibiotic althiomycin ^15^, red pigment prodigiosin and biosurfactant serrawettin W1 ^16^. Here, we aim to combine strengths of genome mining and pangenome analysis to assess the potential of enterobacteria to produce diverse secondary metabolites and understand the association between secondary metabolism and other biological functions (Fig. S1).

First, we set out to examine 2627 complete genomes from 51 different genera of enterobacteria downloaded from the PATRIC database v3.5 ^17^ in order to investigate the number and kinds of secondary metabolic BGCs the different genera might contain (Fig. S2, Data S1). Using antiSMASH v4 ^1^, we detected a total of 8604 BGCs across 2627 genomes with the global average of 3.27 BGCs per genome across enterobacteria (Data S2). *Photorhabdus*, *Xenorhabdus* and *Serratia* are some of the genera with a relatively higher richness in BGCs and have been previously described to produce diverse secondary metabolites^10,15,16^. We observe that there is a genus dependent association between the number of BGCs and the size of the genome (Fig. 1). Interestingly, three of the very small genomes (length < 0.5 Mb) from *Buchnera*, endosymbionts of aphids, also carry a BGC from the aryl polyene class, which are polyketide-derived pigments similar to carotenoids (Fig. 1). The detected BGCs are of 63 different types, inferred through the rule-based antiSMASH BGC detection logic, which includes 28 individual and 35 hybrid BGC types (Table S1). Various BGC types include those encoding for NRPS (2774), thiopeptides (2477), siderophores (693), PKS-NRPS hybrids (614), bacteriocins (612), and 58 other types of BGCs (1446) spread throughout diverse genera (Fig. S3, Fig. 1). A phylogenetic tree of 50 genomes from different genera (Table S2), constructed using all shared proteins and maximum likelihood algorithm (RAxML) ^18^, displayed evolutionary relation of BGCs across enterobacteria (Fig. 1). Most genomes possessed at least one NRPS BGC encoding siderophore (enterobactin or a similar derivative) and one uncharacterized thiopeptide BGC that could be associated with the YcaO superfamily of proteins involved in post-translational modifications of ribosomal protein S12 in *E. coli* ^19^. Enterobacterial genomes are observed to have higher diversity in NRPS type BGCs especially in *Photorhabdus* (avg. 8.4 BGCs), *Xenorhabdus* (avg. 6.6 BGCs) and *Serratia* (avg. 4 BGCs) (Fig. S3). The enterobacteria thus contain a greater richness of BGCs than previously realized.

**Fig. 1.**
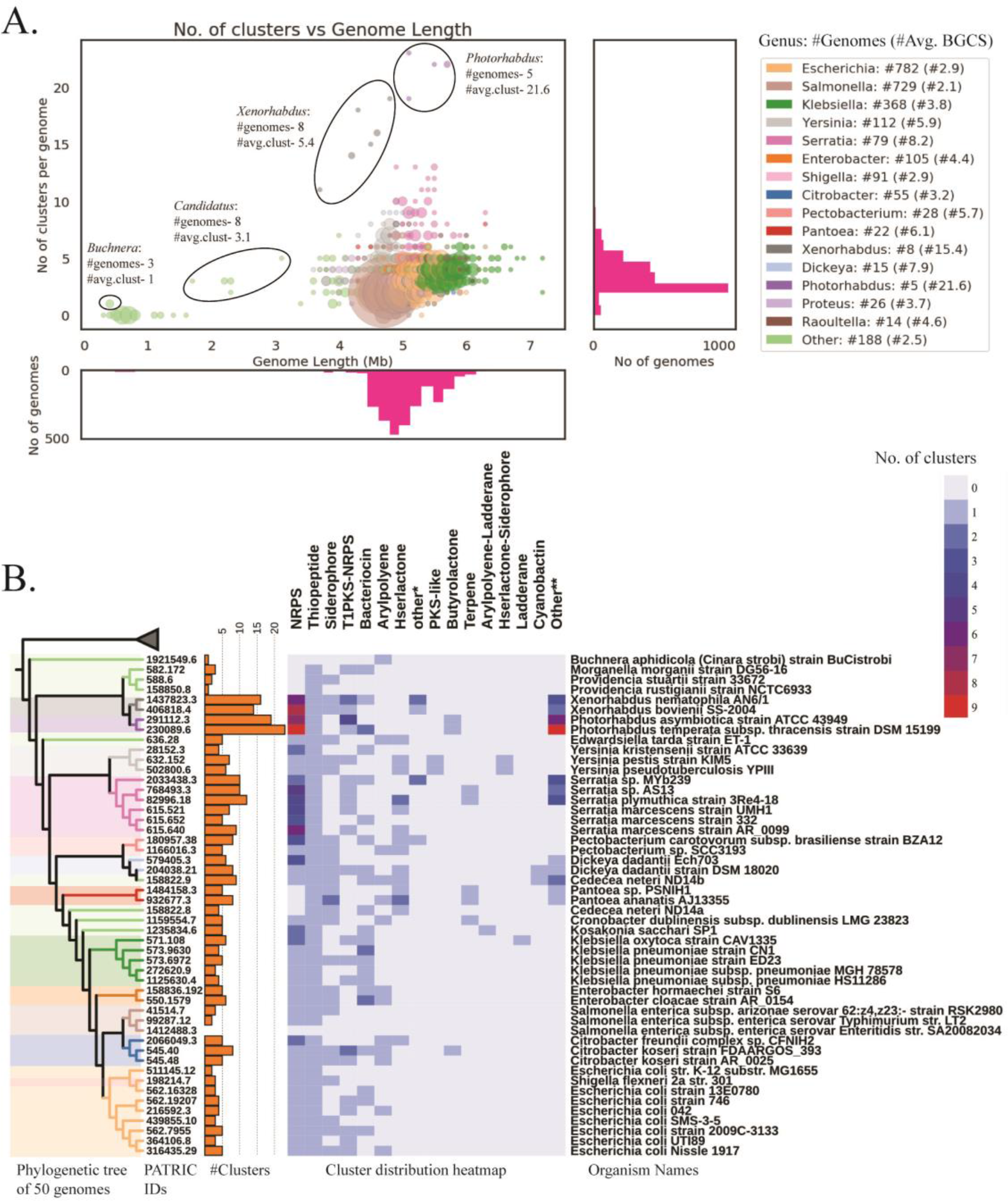
Distribution of BGCs detected across diverse genera of enterobacteria. A. A scatter plot showing the association between the number of BGCs detected across 2627 enterobacterial genomes versus genome length. The size of the circles represents the number of genomes, colors denote major genera. Associated histograms represent the distribution of BGCs in genomes (right) and lengths of genomes (bottom). The number of genomes per genera and average number of BGCs are denoted along with the color legend. B. Distribution of presence of BGCs of various types across 50 selected genomes from 21 diverse genera of enterobacteria. The phylogenetic tree was generated based on all shared proteins and using maximum likelihood algorithm (RAxML) with out-group of 5 genomes from neighboring clades of Gammaproteobacteria (Table S2). The leaves of the tree are colored based on different genera of enterobacteria. Bar chart with the number of BGCs per genome, PATRIC genome accessions and organism names are also mentioned.

Second, we asked how many of the detected BGCs are distinct and how many of them have been assigned to well-characterized secondary metabolites. We used the BiG-SCAPE pipeline ^20^ to compare sequences of 8604 detected BGCs and 1795 known BGCs from the MIBIG database ^21^. BiG-SCAPE uses a weighted combination of domain sequence similarity, Jaccard index, and adjacency index to define a raw distance metric to generate a sequence similarity network. The sequence similarity network suggested 212 distinct families among enterobacterial BGCs (Fig. 2), with 294 BGCs being singletons (Data S3). Surprisingly, only 20 families were associated with previously characterized BGCs from the MIBIG database, whereas the remaining 192 families might encode for novel secondary metabolites (Fig. S4). It is important to note that many of the well-studied secondary metabolites from *Photorhabdus* and *Xenorhabdus* ^*9,10*^ and their associated BGCs are not incorporated in the MIBIG database, thus limiting our prediction capabilities. Adjacency matrices calculated for three of the large families reveal intra-family diversity among BGCs (Fig. S6-S8). It is known that different enterobacteria produce a variety of siderophores derived from enterobactin, such as salmochelin that act as a bacterial evasion mechanism against the mammalian protein siderocalin in *Salmonella* strains and some uropathogenic *E. coli* ^*22*^. Thus, further analysis of large families of BGCs provides avenues to explore the diversity of compounds and their physiological roles.

**Fig. 2.**
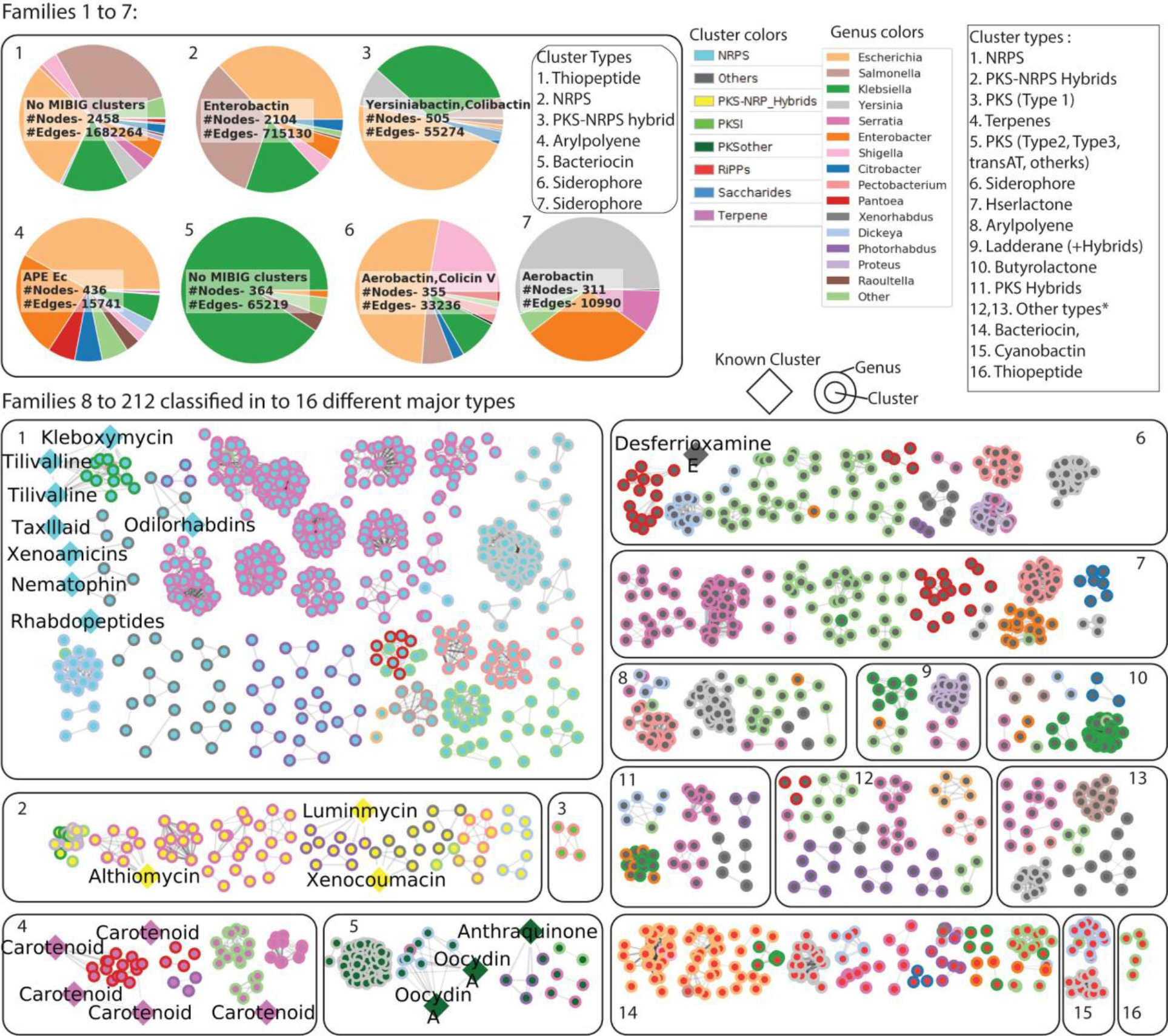
Global network view of BGCs across Enterobacteria. Sequence similarity network of 8604 BGCs with 1795 known BGCs from the MIBIG database reveals 212 distinct families or disconnected components of the network. The largest seven families have large, highly connected networks (Fig. S5) and are represented by pie-charts in A-G. The number of nodes, number of edges, and any known MIBIG BGCs are mentioned in the pie-chart. Different colors and numbers on the perimeter denote different genera and the number of BGCs per genera from that family. For the remaining smaller families, we used Cytoscape to visualize the networks which are grouped according to the class of BGCs in 1-13. Each node represents a biosynthetic gene cluster, where the color of the inner circle denotes the class of the BGC and the color of the outer circle denotes different genera. Edges between two BGCs denote the similarity metric detected using the BiG-SCAPE pipeline. Known BGCs from MIBIG are larger diamond-shaped nodes with known BGCs as labels.

Third, we further examined the genetic structure variations of a particular BGC encoding for colon cancer-related colibactin across diverse genera. The third-largest family of PKS-NRPS BGCs had 503 clusters, where 67 and 181 BGCs are directly similar to colibactin and yersiniabactin respectively and the remaining 255 BGCs were distantly similar (not immediate neighbors in the network) (Fig. 3). Yersiniabactin is observed to be present whenever colibactin is present across genera as also seen earlier ^23^. Yersiniabactin and colibactin clusters are located close to each other in most cases, with the exception of some *Citrobacter sp*. (Fig. 3). The phylogenetic relationship between all of 67 colibactin clusters is further calculated and visualized using CORASON software, which is an integrated part of the BiG-SCAPE pipeline ^20^ (Fig. S9, Data S4). The genus dependent variations in the genetic structure of colibactin BGCs arise mainly due to differences in the intermediate region of colibactin and yersiniabactin core clusters (Fig. 3). Further analysis of the intermediate region between the two core clusters suggested the presence of genes encoding enzymes involved in the Type IV secretion system (T4SS) called VirB in *Klebsiella sp*., *Enterobacter sp*. and *Citrobacter sp.* ^*24*^. Interestingly, no such genes related to secretion systems were observed in the intermediate region for *Escherichia sp*. Neighbouring location of T4SS to colibactin biosynthetic genes in some species hints towards a possible association of this secretion system with an export mechanism of colibactin, which still remains elusive ^25^.

**Fig. 3.**
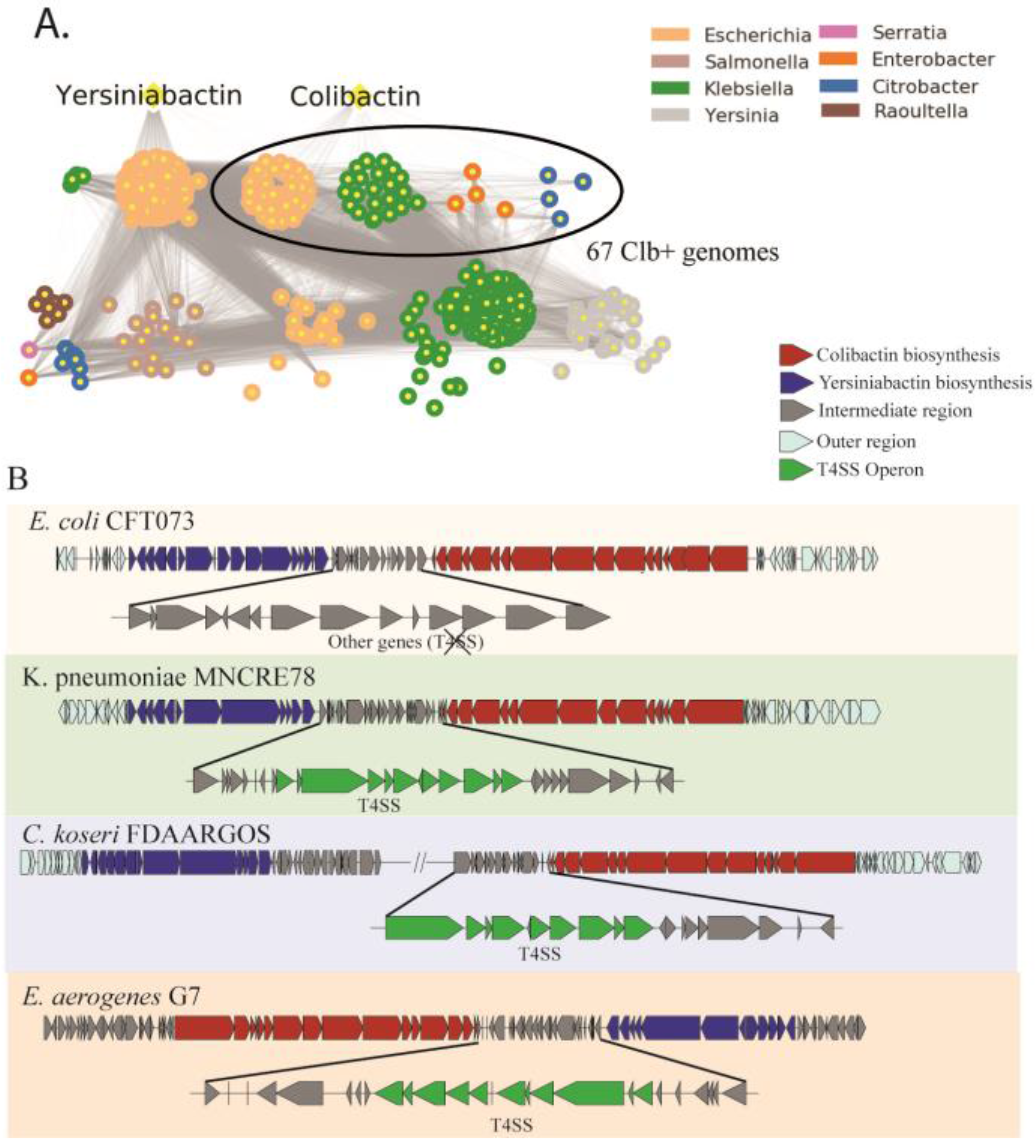
Pangenome reconstruction of 36 PKS containing *Escherichia* genomes followed by comparative genomics. A. Similarity network of PKS-NRPS hybrid clusters from across multiple genera (denoted by colors) encoding yersiniabactin and colibactin. B. Genetic structure of colibactin BGCs across different genera. For *Citrobacter koseri*, the yersiniabactin and colibactin gene clusters occur at different positions in the genomes. The different intermediate regions between the colibactin and yersiniabactin are pointed out.

Fourth, we asked if the comparison of as many as 782 genomes of *Escherichia sp*. from the dataset can guide us to detect specific genetic and functional associations with the presence of colibactin. Here, we investigated *Escherichia* genomes possessing colibactin using pangenome reconstruction and genome-wide association techniques ^5,26,27^. For 36 colibactin-containing *Escherichia* genomes, a total of 14,017 different genes were detected in the pan-genome (total genes found across genomes), which was reconstructed using Roary ^28^(Table S3, Data S5). The core-genome (genes shared among genomes) was composed of 2946 genes. Conversely, 4407 genes were uniquely present in only one of the genomes (Fig. 4). We compared sequences of 2937 genes (remaining 9 were tRNA genes) from the reconstructed core-genome against all 782 *Escherichia* genomes present in the dataset using bidirectional best blastp hits (Fig. 4, Data S6). Examining the gene presence/absence heatmap (Fig. 4) revealed a set of 110 genes that are present in all colibactin containing genomes and are absent in more than 90% of the genomes missing both colibactin and yersiniabactin (Table S4). This set of colibactin associated genes are spread across the genome describing various biological functions including secondary metabolism, secretion system, amino acid metabolism, fimbrial usher chaperone pathways, regulation among others. Thus, the growing availability of complete gene sequences enables us to define a gene set that is associated with a particular BGC.

**Fig. 4.**
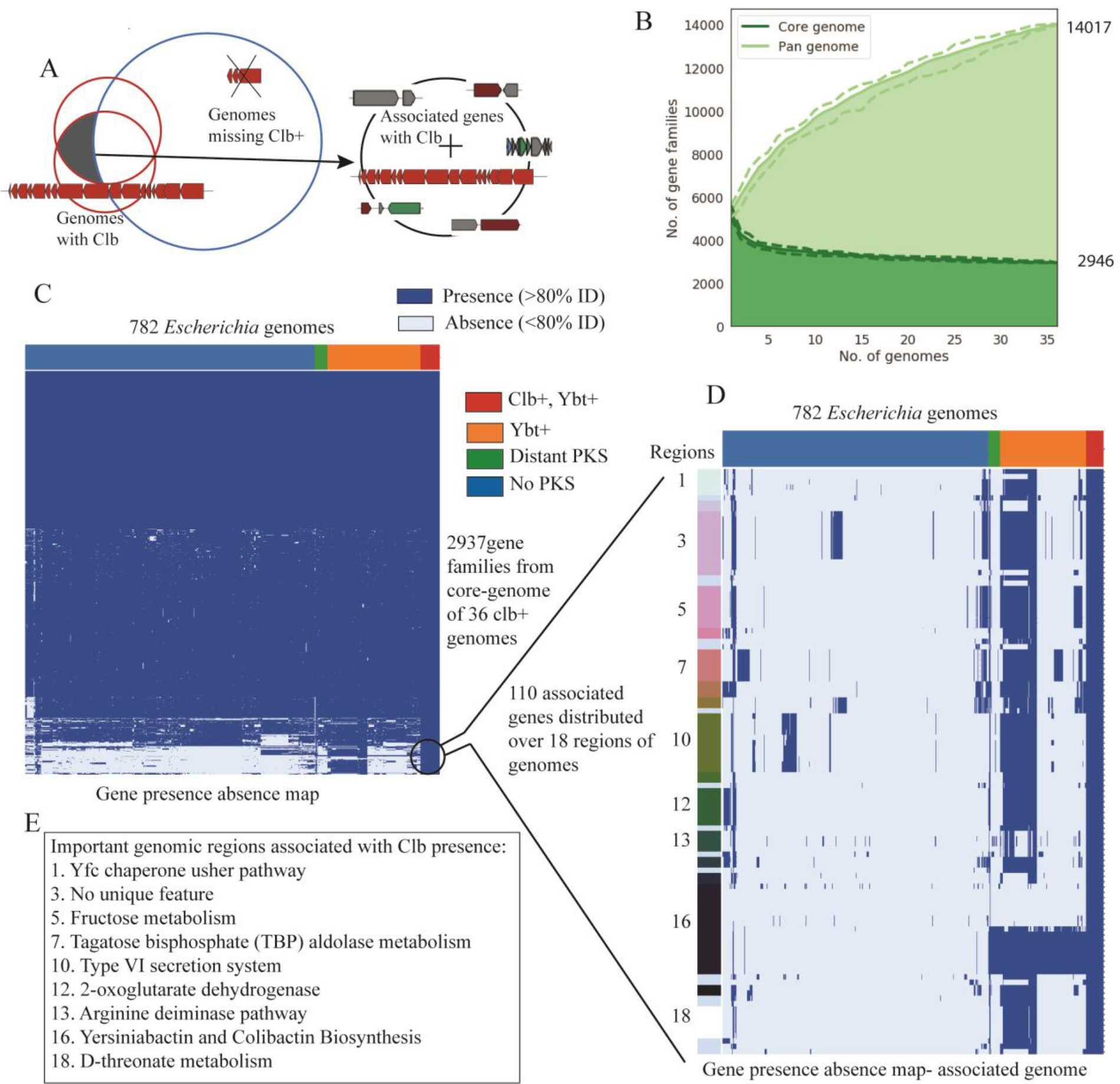
Phylogenetically associated genetic signatures with the presence of colibactin BGC. A. Schema describing comparing core-genome of colibactin containing genomes against those missing colibactin to detect a set of genes associated with presence of colibactin. B. 36 genomes possessing colibactin PKS island are selected for the reconstruction of the core-genome (common across all) and pan-genome (total genes). C. Gene presence/absence heatmap of 2937 core genes (9 tRNA genes are ignored from 2946) as compared against all 782 genomes of *Escherichia*. Columns represent genomes divided into four groups: i) both colibactin and yersiniabactin clusters (36); ii) only yersiniabactin cluster (176); iii) PKS-NRPS clusters that are distantly similar to yersiniabactin or colibactin (24); and iv) none of the PKS clusters (546) (Fig. 3). Here, the gene presence is defined if the homolog identity was higher than 80%. D. Heatmap of 110 genes that are present in all 36 genomes and are absent in more than 90% of PKS absent *Escherichia* genomes. The rows are grouped by colors depending on the presence of genes in different regions/ operons in the genome. The few selected regions are further described (Table 4). E. List of important genomic regions from an associated set of genes with common biological functions.

Finally, we investigated some of the genes that were observed to be highly conserved across colibactin-containing genomes. In total, 94 of the 110 genes were observed located in 18 different genomic regions, and 14 genes were located alone throughout the genome (Table S4). The largest continuous region of in associated gene sets encoded for colibactin (15 genes) and yersiniabactin (12 genes) biosynthesis. Most interestingly, one region of 11 genes encoded for the assembly of Type VI Secretion System (T6SS), whereas genes encoding similar T4SS were shown to be present in the neighborhood of clusters in other genera than *Escherichia* (Fig. 3). Both T4SS and T6SS contain periplasm-spanning channels and secrete proteins from the cytoplasm outside the cell and across an additional host cell membrane, delivering secreted substrates directly to the cytosol of a target cell ^29^. Thus these secretion systems provide an efficient way to inject substrates in a contact-dependent manner, which is required for genotoxic effects of colibactin as discovered earlier ^30,31^. Some of the associated genes are also conserved in part of the Yersiniabactin possessing genomes (variable from 0 to 57%) revealing the importance of the associated functions during evolution for PKS-NRPS clusters in *Escherichia* (Table S4). The identified set of associated genes suggests functions with putative roles in the overall function of a BGC.

As the number of genomes being sequenced is growing rapidly, the genome mining approaches can give new insights in predicting enterobacterial strains with the potential to produce novel compounds as well as identify novel phylogenetically associated functions with these compounds. Given that enterobacteria are studied extensively and the role of secondary metabolites in human physiology, it is surprising that most of the families of BGCs detected here have not yet been functionally studied. This finding motivates a strong need to further investigate enterobacterial secondary metabolism, which can open up the possibility to discover novel secondary metabolites and their physiological roles. The predicted associated set of genes of the colibactin BGC will open up many avenues to explore the mechanisms related to the activity of colibactin in pathogenic *E. coli*. Given the extensive knowledge of primary metabolism and state-of-the-art metabolic engineering tools available for *E. coli*, strategies to understand secondary metabolism and associated functions can provide a guide to design host strains to efficiently express heterologous clusters from other bacteria.

## Methods

### Collection and quality assessment of enterobacterial genomes

We collected 2139 genome entries from PATRIC database ^17^ of Enterobacteriales which are annotated as ‘complete’. The quality of genome assembly is important for better prediction of genome mining as well as for the pangenome analysis. To curate the data, we manually removed 12 genome entries due to irregular genome size, a high number of contigs, or low N50 score (Fig. S1, Data S1). The distribution of genome size against GC content showed that there are very few genomes that are outliers when compared to the average genome size of a particular genus (Fig. S1). Genomes from other genera including endosymbiont genera such as *Buchnera*, which have relatively lower genome sizes, are not removed from the analysis. We note that 1135 genomes had a single contig and 1424 genomes had N50 score higher than 90% of the genome size suggesting the high quality of assemblies. To understand the redundancy in the genomic datasets, we note that 37 organisms had more than one genome sequences in the dataset (Data S1). For example, well studies strain *E. coli* str. K-12 substr. MG1655 has 11 genome sequences in the dataset. As the number of such redundant genomes is relatively less, we considered all these genomes in our final dataset. Thus, the final dataset contained 2127 genomes spread across 51 different genera with the majority belonging to *Escherichia*, *Salmonella*, and *Klebsiella* (Data S1).

### Genome mining of secondary metabolites BGCs across enterobacteria

The antiSMASH v4 ^1^ was used for genome mining of secondary metabolite BGCs in 2627 enterobacterial genomes. We used ‘.gff’ files downloaded from the PATRIC database for gene annotations with the antiSMASH run. As all subsequent steps require complete BGCs, we filtered out 99 BGCs that are detected on the contig edges (Data S2). The final data set contains 8604 BGCs detected across genomes of enterobacteria (Data S2). For genus-specific analysis and visualization, we considered the top 15 genera with the highest total number of clusters per genus. The remaining 36 genera are all classified as ‘Other’. Similarly, we considered the top 15 most occurring types of the BGCs as defined in antiSMASH output and classify the remaining 48 BGC types as ‘Other’ (Table S1). The distribution of the average number of BGCs of each type across genera is visualized in the heatmap (Fig. S3).

### Phylogenetic tree of selected enterobacterial genomes with different BGC distributions

A set of 50 genomes was selected from various genera with different distributions of BGC types as detected through genome mining (Table S2). A phylogenetic tree was constructed with PATRIC ^17^ services for tree building based on all shared proteins and using maximum likelihood algorithm (RAxML) ^18^. The out-group set used for the tree construction involved five genomes of neighboring clades in Gammaproteobacteria from the PATRIC database, which are 1) *Mannheimia succiniciproducens* MBEL55E; 2) *Pasteurella multocida* subsp. multocida str. Pm70; 3) *Photobacterium profundum* SS9; 4) *Vibrio fischeri* ES114; and 5) *Vibrio cholerae* O1 biovar El Tor str. N16961. The phylogenetic tree was visualized using iTOL v4(Interactive Tree of Life) ^32^, with additional panels representing a bar chart of total BGC distribution and heatmap of BGC type distribution across all genomes (Fig 1).

### Sequence-based similarity network of BGCs detected in Enterobacteria

BiG-SCAPE ^20^ software was used to generate a sequence-based similarity network of 8604 detected clusters. Here, ‘*--mibig’* parameter was used for comparison against 1795 known BGCs from the MIBIG database ^21^. For the similarity index between any two clusters, we used *raw_index* metric from BiG-SCAPE output. We tried 3 different cutoffs of 0.3, 0.5, 0.7 on *raw_index* to define the similarity network. As the sequences of many BGCs are very similar to each other, we used a cutoff of 0.3 for the final network (Data S3). The comparison was carried out separately for the major classes of BGCs by using the parameter *--hybrids-off*.

### Detecting families of BGCs from sequence-based similarity network

The distinct connected components, based on a cutoff of 0.3 on *raw_index*, are denoted as distinct families of BGCs. Using the similarity network, 212 completely distinct families were detected. The distribution of BGC families across genomes of different genera is represented by a heatmap (Fig. S4). As some of the BGC families are highly frequent than others in enterobacteria, the top 7 largest families are visualized using pie-charts (Fig. 2). The networks for remaining smaller families are visualized using Cytoscape ^33^ and are categorized based on BGC classes (Fig. 2). For the top 7 largest families, the network visualization using Cytoscape can be found in Fig. S5. Further, we analyzed the top 3 families by calculating the adjacency matrix of the network to understand the intra-family diversity in BGC contents (Fig. S6-S8). For family 2, enterobactin was manually assigned to BGCs from *E. coli* K-12 MG1655 strain because no similar BGC was detected in MIBIG standard database. We observe that the MIBIG database does not have an entry for a well-known enterobactin BGC from *E. coli*. BGCs similar to known colibactin clusters from different genera are compared using BiG-SCAPE to 3 identity subfamilies describing diversity in the genetic structure of BGCs across genera (Fig. S9, Data S4). Genetic structure of the four of the BGCs from different genera, 199310.4.cluster004 (*Escherichia coli* CFT073), 573.7238.cluster004 (*Klebsiella pneumoniae* MNCRE78), 545.40.cluster006 (*Citrobacter koseri* FDAARGOS_393) and 548.155.cluster001 (*Enterobacter aerogenes* G7), are further analyzed (Fig. 3).

### Characterization of colibactin associated gene sets for *Escherichia*

We selected 36 genomes of *Escherichia sp*. possessing the colibactin gene cluster for pangenome analysis. Pangenome reconstruction was carried out using Roary software ^28^ (Fig. 4, Data S5). We selected genes from the core genome that are commonly present across 36 colibactin cluster-possessing genomes for further investigation. To analyze the specific functions associated with colibactin BGC, we compared the 2937 genes coding for proteins from the core genome against all of the 782 genomes using bi-direction best hits using diamond (Data S6). The gene presence-absence heatmap was generated with genes with greater than 80% identity defined to be similar (Fig. 4). The set of 110 genes that are present across all of colibactin containing genomes and missing in more than 90% of no pks containing genomes are further investigated for their putative role in the function of colibactin BGC (Fig. 4, Table S4). The set of associated genes that appear next to each other in the genome is defined as a genomic region. In total, 18 such genomic regions can be defined incorporating 94 of the 110 associated genes. Many of the regions with genes encoding for a pathway or cluster of genes having a similar functional roles are assigned a common putative function that can be associated with the role of colibactin.

## Data availability

The genome sequences used in this study are available at the PATRIC database with accessions mentioned in supplementary materials. All other data is available in the main text or supplementary materials.

## Code availability

For genome mining and generating similarity network of biosynthetic gene clusters, we used antiSMASH v4 and BiG-SCAPE respectively. For pangenome analysis, the Roary pipeline was used. Custom Jupyter notebooks used for data analysis and visualization are uploaded at https://github.com/OmkarSaMo/Pangenome_Sec_Met.

## Acknowledgments

We thank Kai Blin and Simon Shaw for helpful discussions and critical advice on the genome mining workflow. We thank Marc Abrams for his comments on the draft. This work was supported by grants from the Novo Nordisk Foundation (NNF10CC1016517, NNF16OC0021746).

## Author contributions

B.O.P and T.W. conceived the study; O.M., T.W. and B.O.P. designed the research; O.M. collected data, performed research, analyzed and interpreted the data, with assistance from C.J.L and J.M; O.M., B.O.P., T.W. drafted the article; O.M., C.J.L., J.M.M., T.W., and B.O.P. critically revised and approved the article.

## Supplementary information

**Fig. S1:**
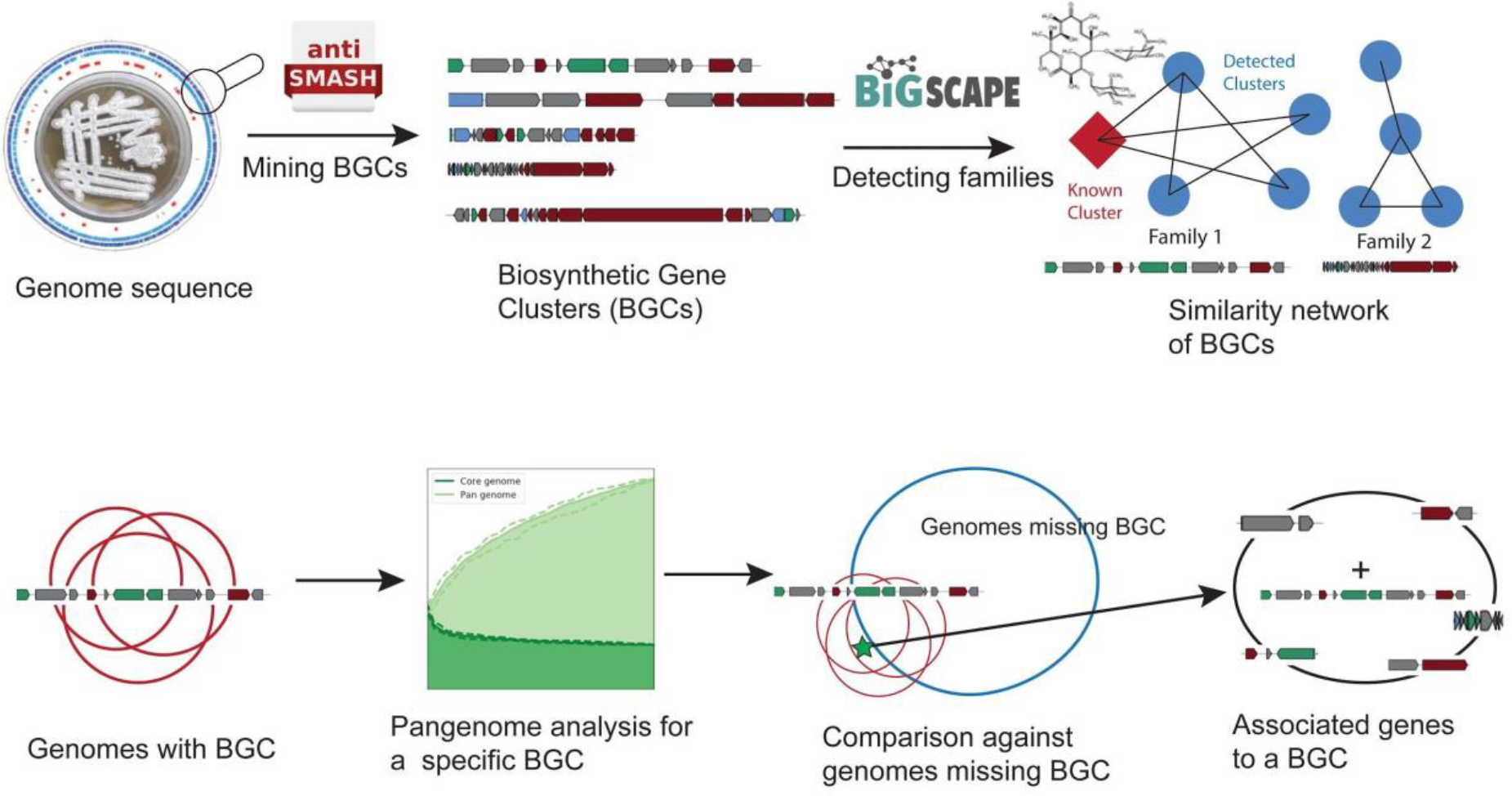
Schematic representation of the genome mining and pangenome analysis workflow. Large set of genomes were mined for BGCs encoding for secondary metabolite biosynthesis using antiSMASH ^1,2^. The diversity of detected clusters and identification of known BGCs was examined using BiG-SCAPE ^20,21^ and MIBIG database ^21^. Further, pangenome analysis was carried out for a set of genomes with having same BGC. The core-genome of genomes with particular BGC was compared against other genomes missing that BGC to identify a set of genes that might have a functional association with the presence of BGC.

**Fig. S2.**
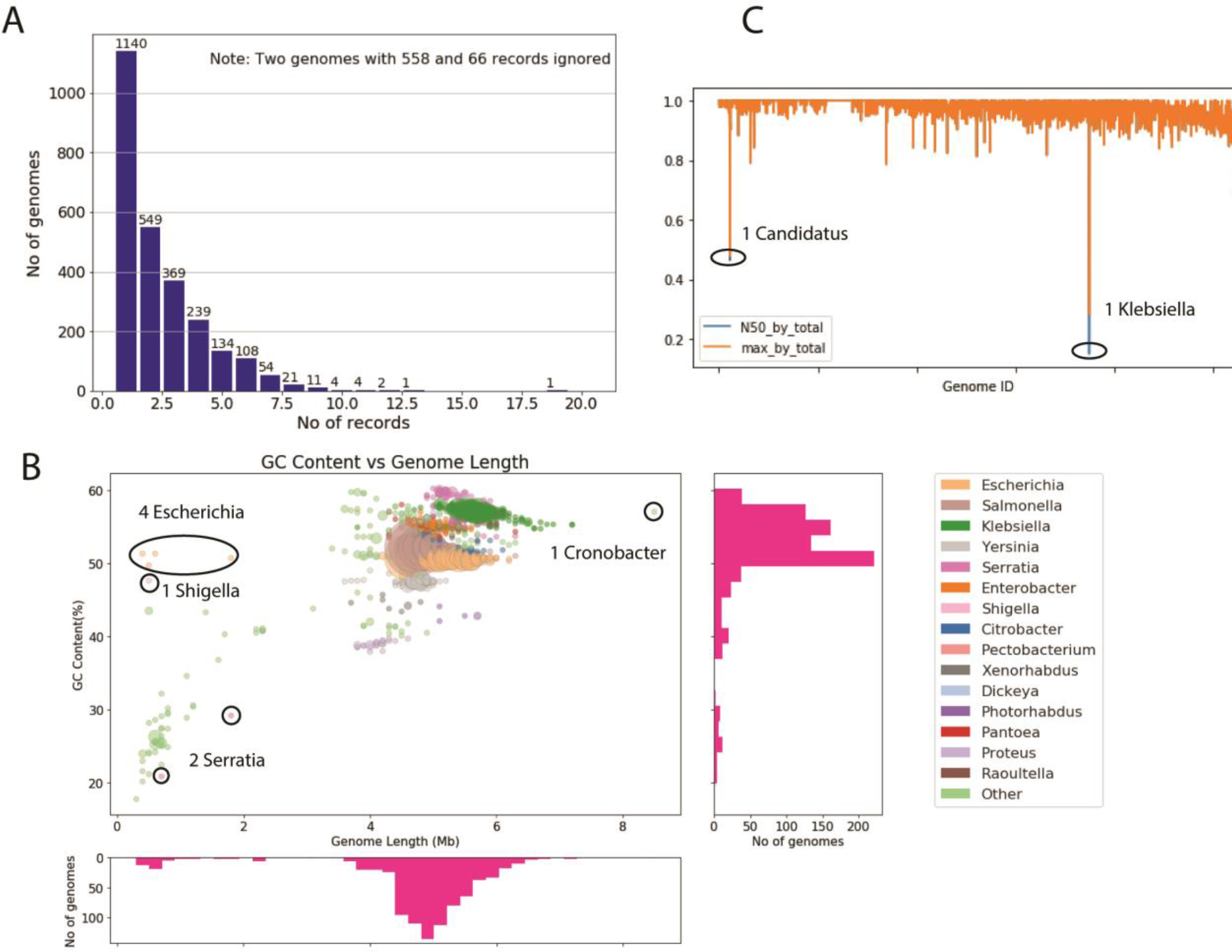
Genome quality assessment and curation of the input dataset. A. Distribution of the number of records per genome suggesting high-quality assemblies. Here, two genomes from the initial set of 2639 are removed due to the higher number of contigs. B. Scatter plot showing the distribution of genome length and GC content of 2637. The size of the circles represents the number of genomes, colors denote major genera. Associated histograms represent the distribution of BGCs in genomes (right) and lengths of genomes (bottom). Highlighted 4 genomes of *Escherichia*, 2 genomes of *Serratia*, 1 genome of *Shigella* and 1 genome of *Cronobacter* are removed from the dataset due to their atypical genome size. C. N50 score analysis showed that the genome assemblies are of high quality. N50 genome length and maximum contig length divided by total length are plotted for the remaining 2629 genomes. Two genomes with low N50 scores are removed to get a final dataset of 2627 genomes for further analysis.

**Fig. S3:**
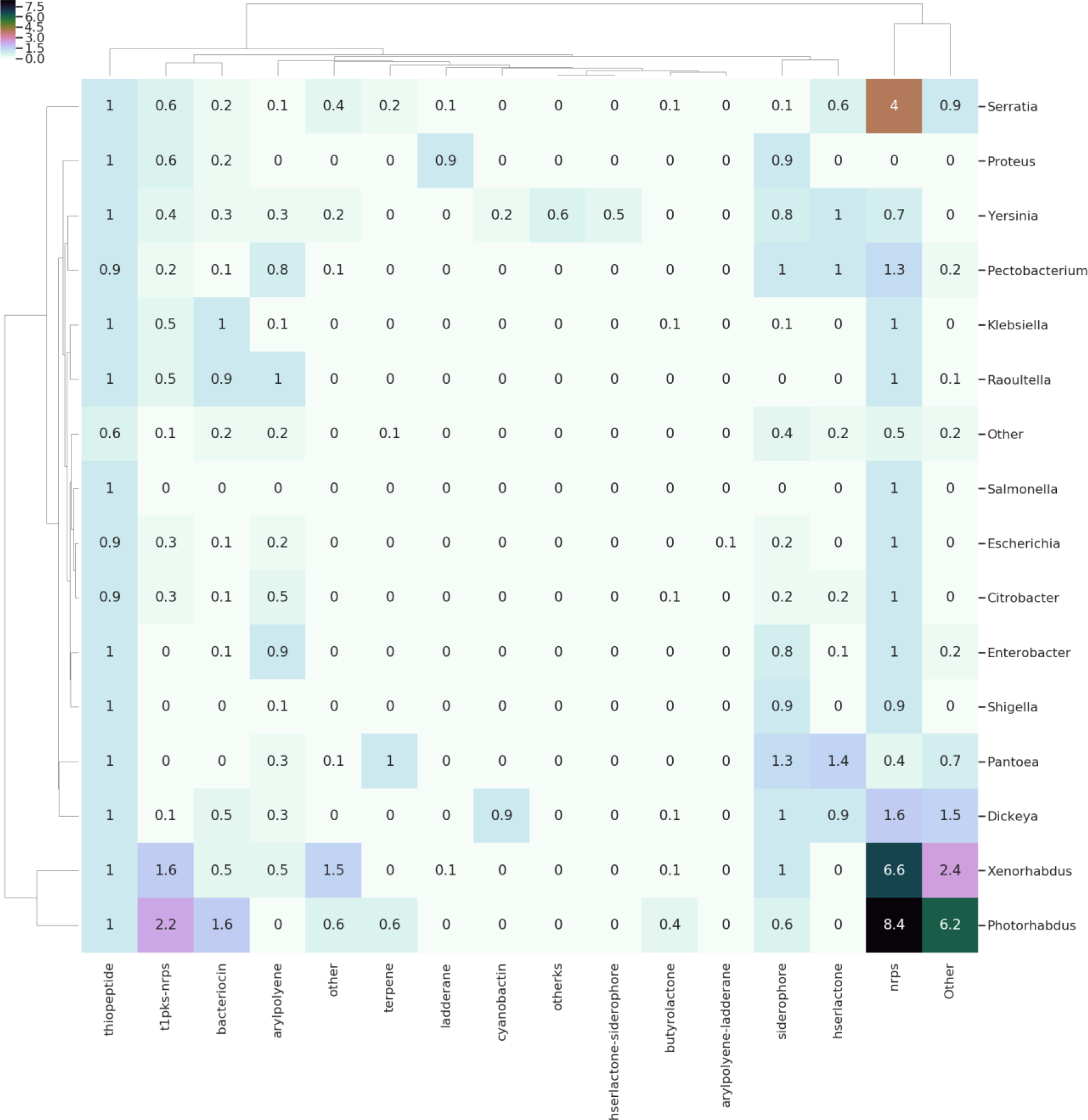
Average number of BGCs of different types across genera. Distribution of the average number of clusters of a particular class (X-axis) detected in the genomes of particular genera (Y-axis).

**Fig. S4:**
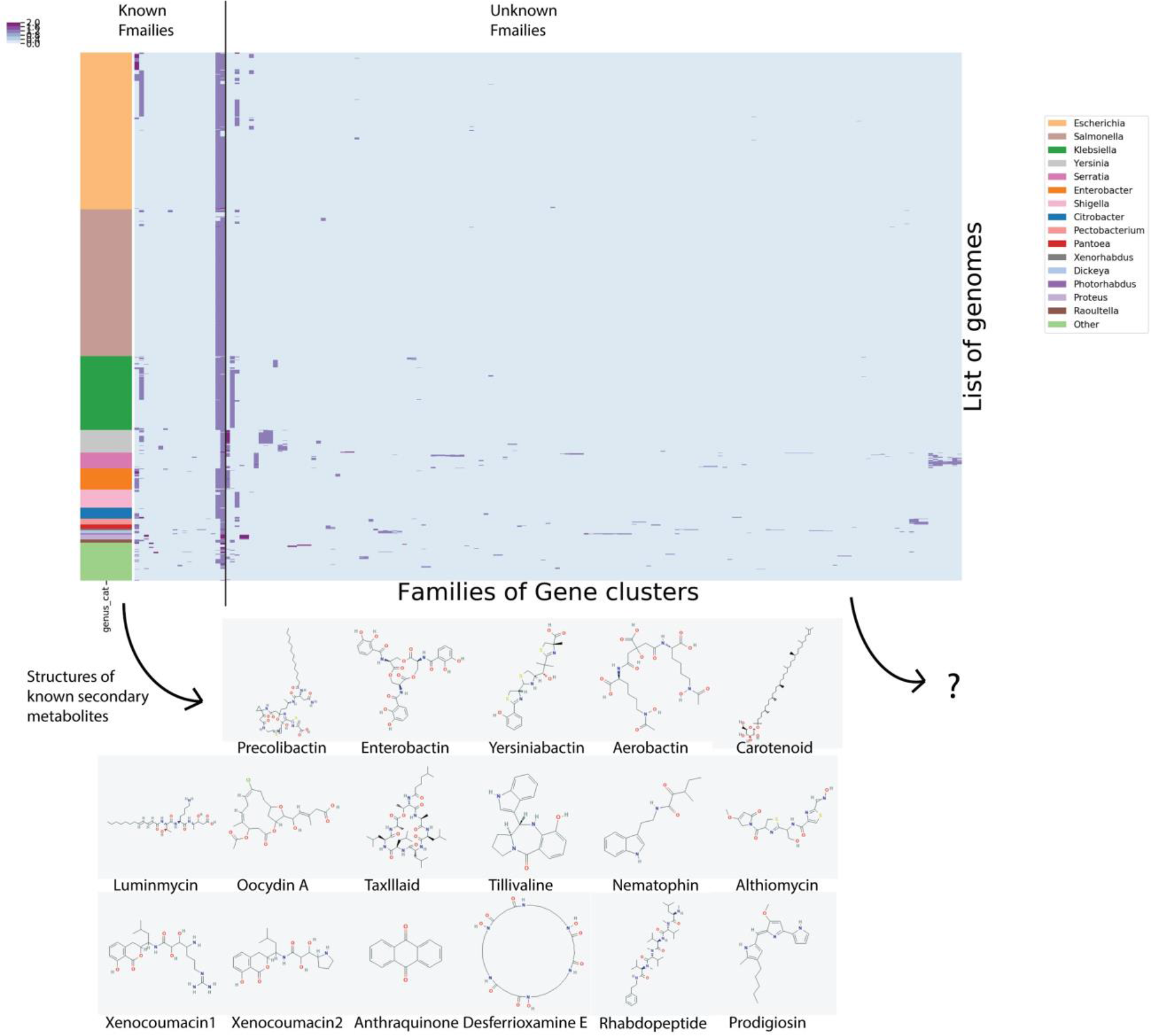
Distribution of BGC families across genomes. A. Distribution of families of BGCs across genomes from different genera. The rows of heatmap represent 2627 genomes ordered according to the genus they belong to. The columns represent the presence of a BGC from one of the 212 distinct families. The columns are split into families that have been associated with known BGCs from MIBIG and others which are new families detected in this study. B. The structures of some secondary metabolites encoded by known BGCs.

**Fig. S5:**
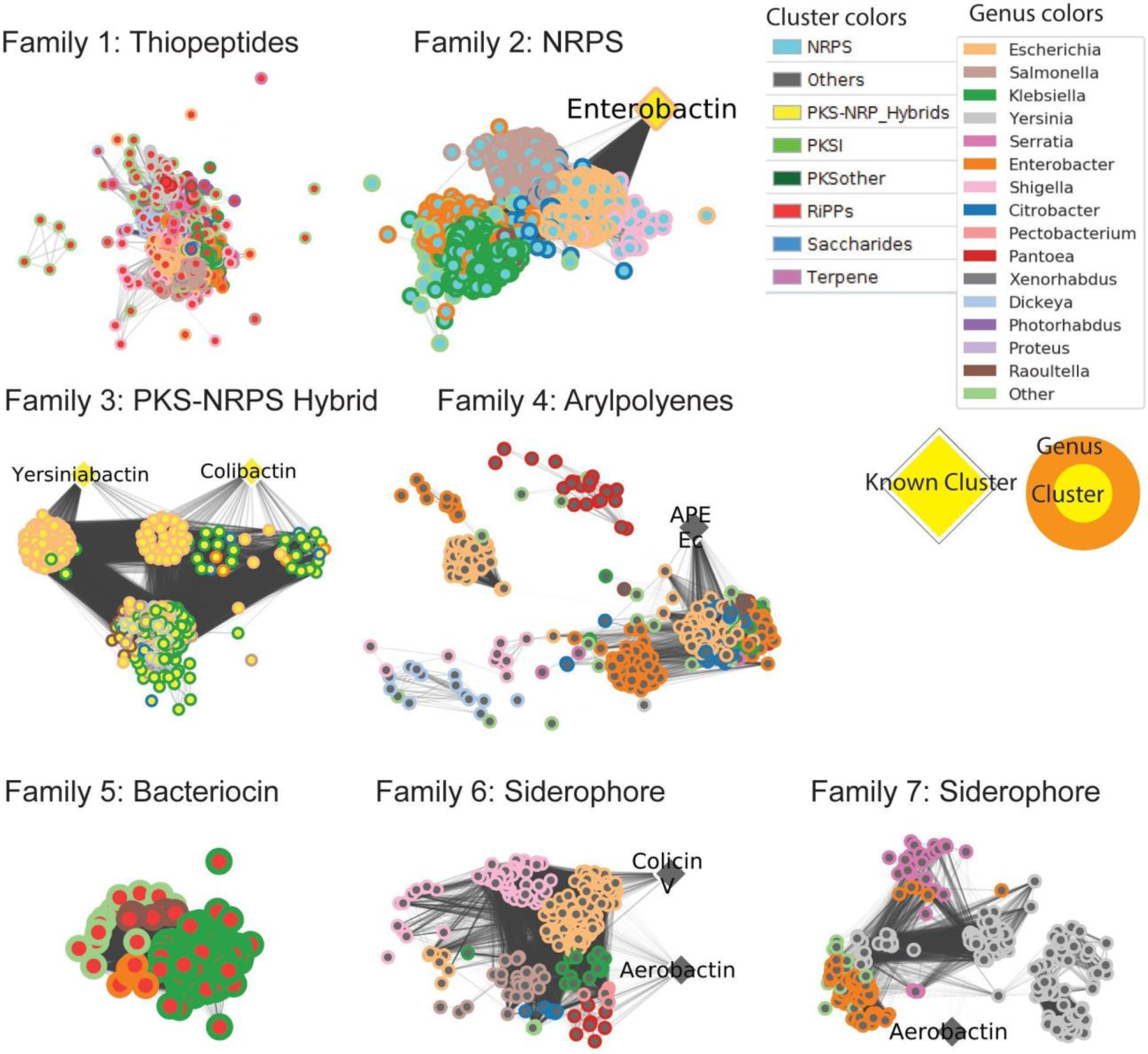
Extended visualization of similarity networks of top 7 largest families of BGCs. Similarity network visualized using Cytoscape for the top 7 largest families. Circular nodes represent BGCs of a particular BGC type (inner circle color) from a genome of a particular genus (outer circle), whereas diamond shaped nodes represent known BGCs from the MIBIG database. The edges represent the similarity between the BGCs calculated using BiG-SCAPE. For family 2, enterobactin was manually assigned to BGCs from *E. coli* K-12 MG1655 strain because no similar BGC was detected in MIBIG standard database. We observe that the MIBIG database does not have an entry for a well-known enterobactin BGC from *E. coli*.

**Fig. S6:**
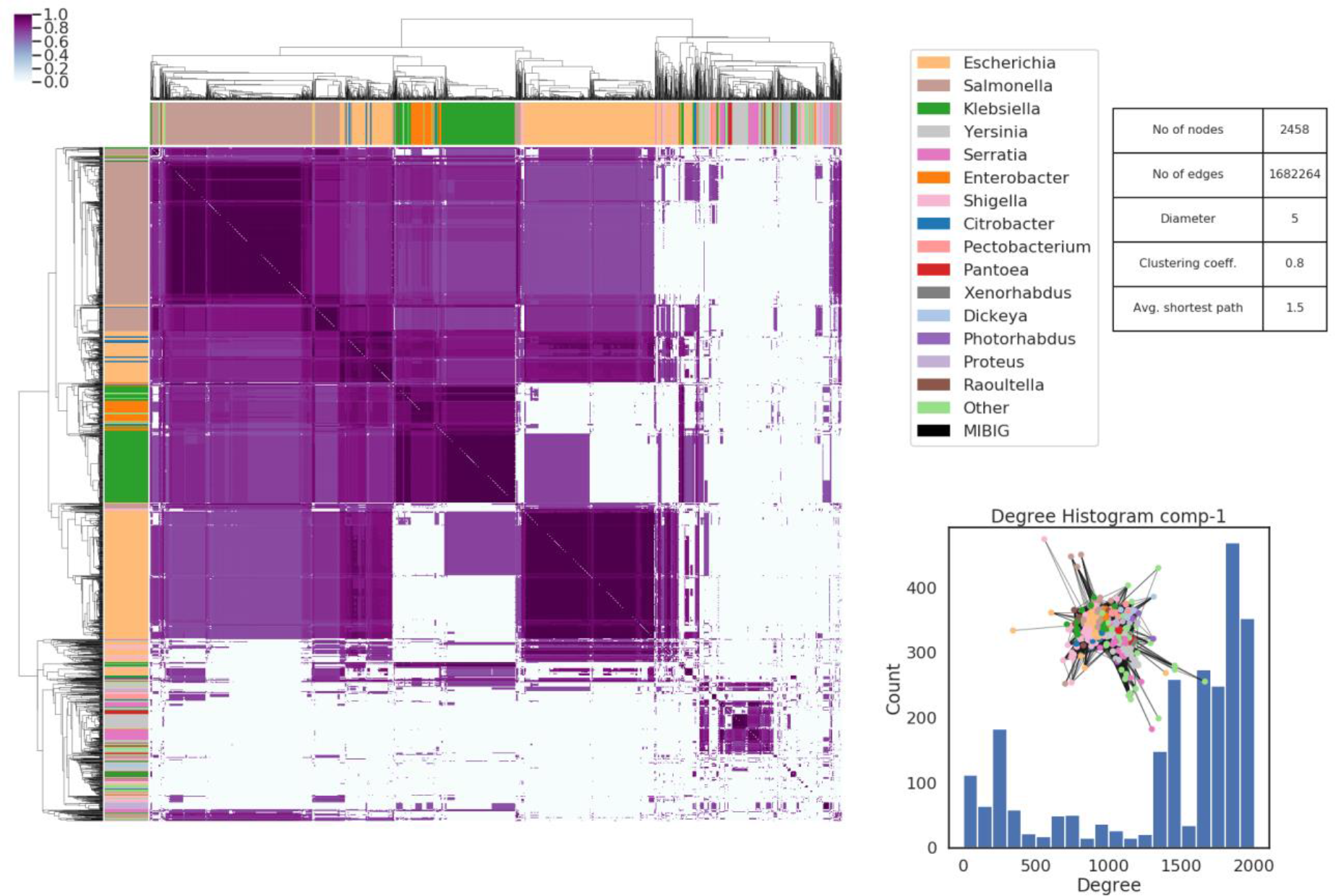
Adjacency matrix for family1 of uncharacterized thiopeptides. The adjacency matrix of the similarity network of family 1. The row and column colors of heatmap represent the different genera of the genome with this BGC. Overview of the network statistics in the table and degree histogram of the network. Note that a cutoff of 0.3 was used for generating similarity networks this the nodes with distance above 0.3 are considered disjoint.

**Fig. S7:**
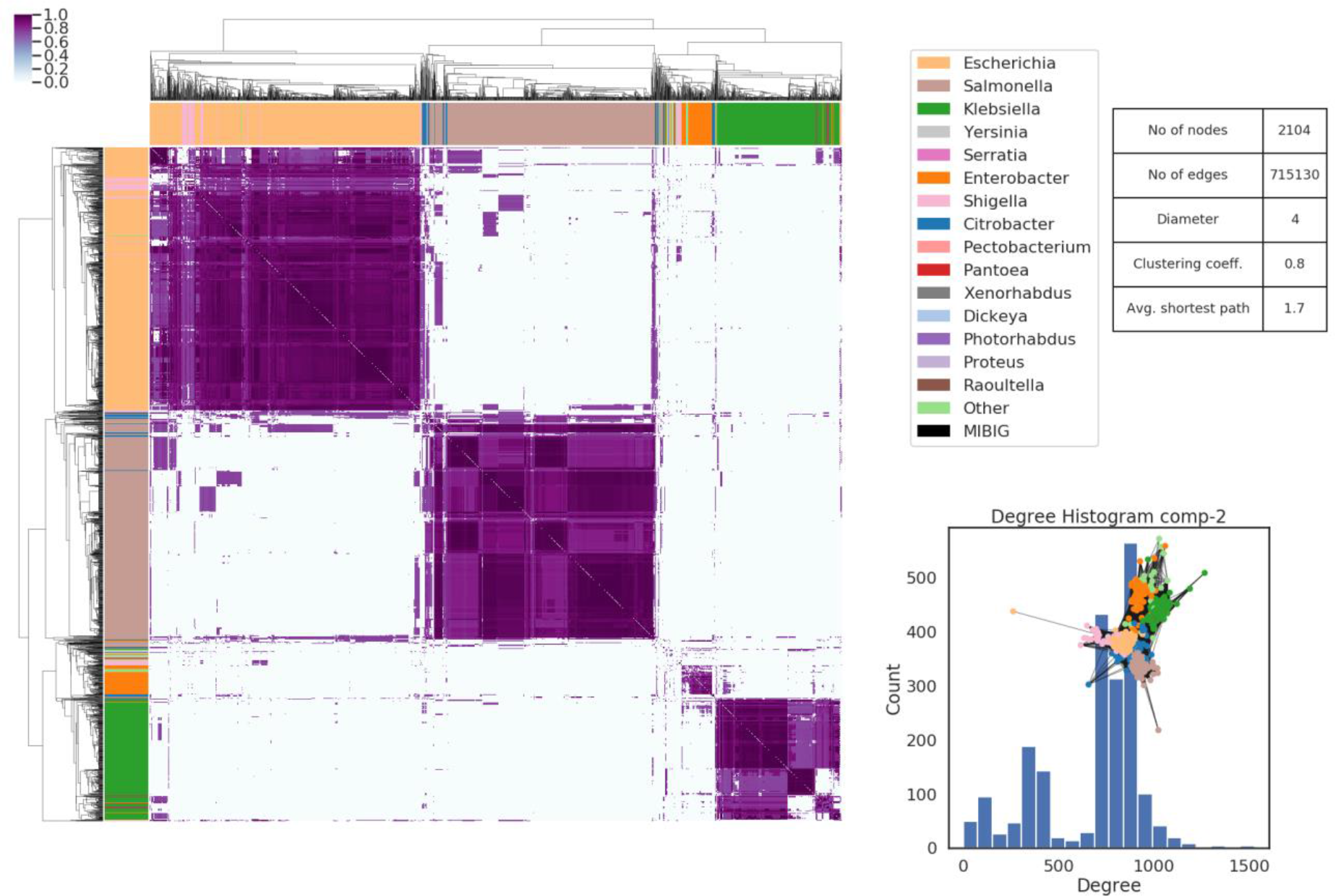
Adjacency matrix for family2 of NRPS encoding for enterobactin and similar derivatives. The adjacency matrix of the similarity network of family 2. The row and column colors of heatmap represent the different genera of the genome with this BGC. Overview of the network statistics in the table and degree histogram of the network. Note that a cutoff of 0.3 was used for generating similarity networks this the nodes with distance above 0.3 are considered disjoint.

**Fig. S8:**
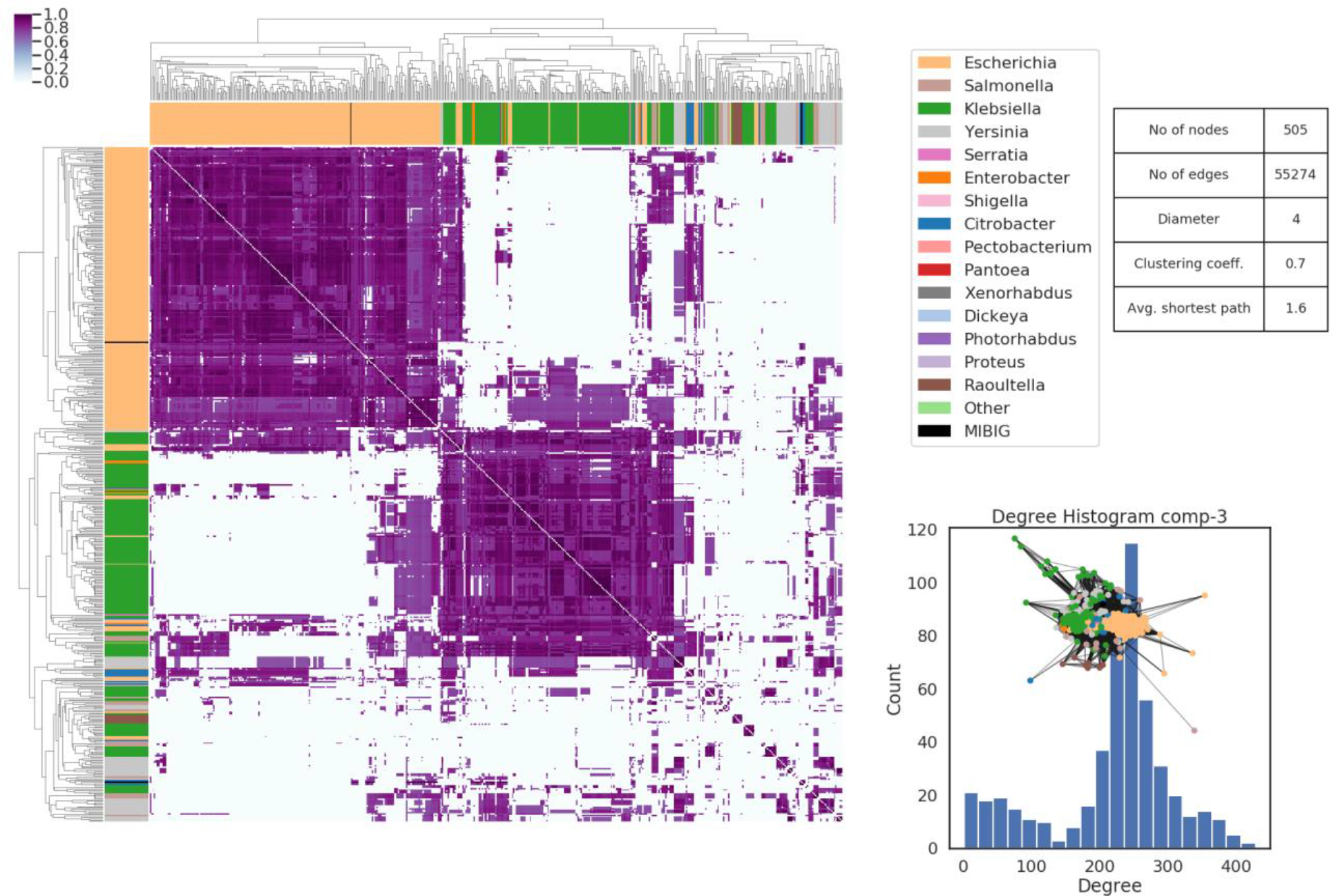
Adjacency matrix for family 3 of PKS-NRPS hybrid BGCs annotated with yersiniabactin and colibactin. The adjacency matrix of the similarity network of family 3. The row and column colors of heatmap represent the different genera of the genome with this BGC. Overview of the network statistics in the table and degree histogram of the network. Note that a cutoff of 0.3 was used for generating similarity networks this the nodes with distance above 0.3 are considered disjoint.

**Fig. S9.**
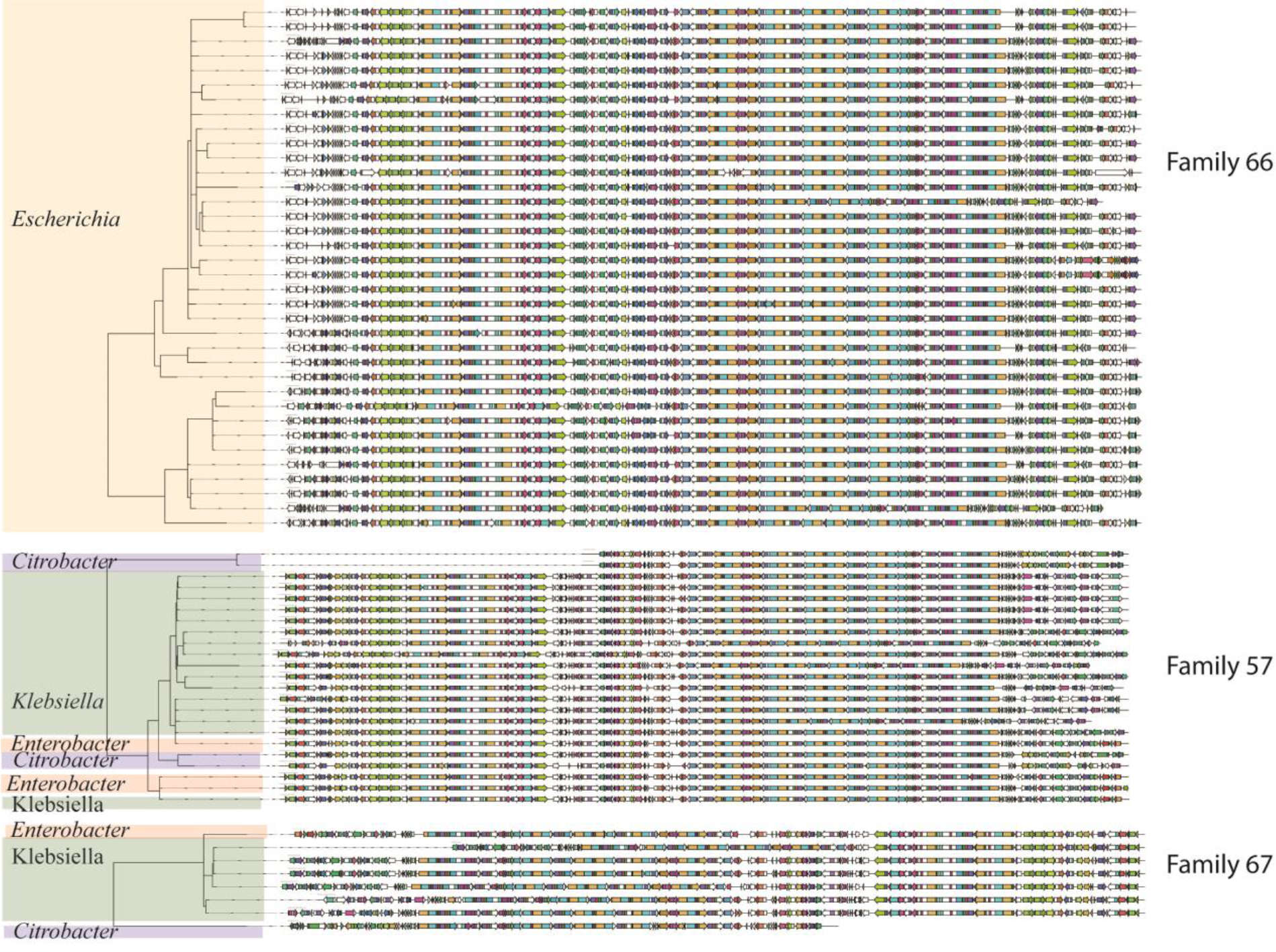
Colibactin BGCs genetic structure distribution. Alignment of genes in colibactin BGC across 67 genomes of different genera, of which 4 are displayed in Fig. 3. The colors on the arrow represent protein domains detected in BiGSCAPE analysis. The phylogenetic tree was generated using CORASON software that is part of BiGSCAPE. The three subfamilies are detected using BiG-SCAPE (Data S4).

**Table S1:** Number of BGCs of different types as defined by antiSMASH rule-based detection logic.

| BGC Type | #BGCs | BGC Type | #BGCs | BGC Type | #BGCs |
| --- | --- | --- | --- | --- | --- |
| **nrps** | **2774** | hserlactone-arylpolyene | 9 | lassopeptide | 2 |
| **thiopeptide** | **2477** | otherks-t1pks-PUFA-nrps | 9 | nrps-siderophore | 2 |
| **siderophore** | **693** | transatpks-otherks | 9 | oligosaccharide | 2 |
| **t1pks-nrps** | **614** | ppysks | 8 | phosphonate-butyrolactone | 2 |
| **bacteriocin** | **612** | transatpks-nrps | 8 | transatpks-otherks-nrps | 2 |
| **arylpolyene** | **418** | nrps-t1pks-PUFA-otherks | 7 | transatpks-t1pks | 2 |
| **hserlactone** | **295** | resorcinol | 7 | bacteriocin-nrps | 1 |
| **other\*** | **95** | nucleoside | 6 | bacteriocin-otherks-PUFA-nrps | 1 |
| **otherks (PKS-like)** | **69** | t3pks | 6 | bacteriocin-phosphonate-butyrolactone | 1 |
| **butyrolactone** | **68** | thiopeptide-nrps | 6 | ectoine-butyrolactone | 1 |
| **terpene** | **61** | phosphoglycolipid | 5 | hserlactone-ladderane-t2pks | 1 |
| **arylpolyene-ladderane** | **56** | t2pks | 5 | hserlactone-nrps | 1 |
| **hserlactone-siderophore** | **52** | ectoine | 3 | ladderane-t1pks-nrps | 1 |
| **ladderane** | **49** | microcin-lassopeptide | 3 | nrps-t1pks-siderophore | 1 |
| **cyanobactin** | **34** | PUFA-otherks | 3 | nucleoside-nrps | 1 |
| t2pks-ladderane | 34 | siderophore-nrps | 3 | phosphonate-nrps | 1 |
| arylpolyene-siderophore | 31 | acyl_amino_acids | 2 | ppysks-nrps | 1 |
| phosphonate | 19 | amglyccycl | 2 | t1pks-butyrolactone-nrps | 1 |
| t1pks | 11 | arylpolyene-nrps | 2 | t1pks-lassopeptide-nrps | 1 |
| blactam | 10 | bacteriocin-siderophore | 2 | t2pks-hserlactone-ladderane | 1 |
| phenazine | 10 | lantipeptide | 2 | transatpks-t1pks-nrps | 1 |

\*Please note here ‘other’ represents a BGC type (not to be confused with ‘Other’ types defined for Fig. 1) that is not assigned to well-known BGCs as per antiSMASH BGC detection logic.

**Table S2:**
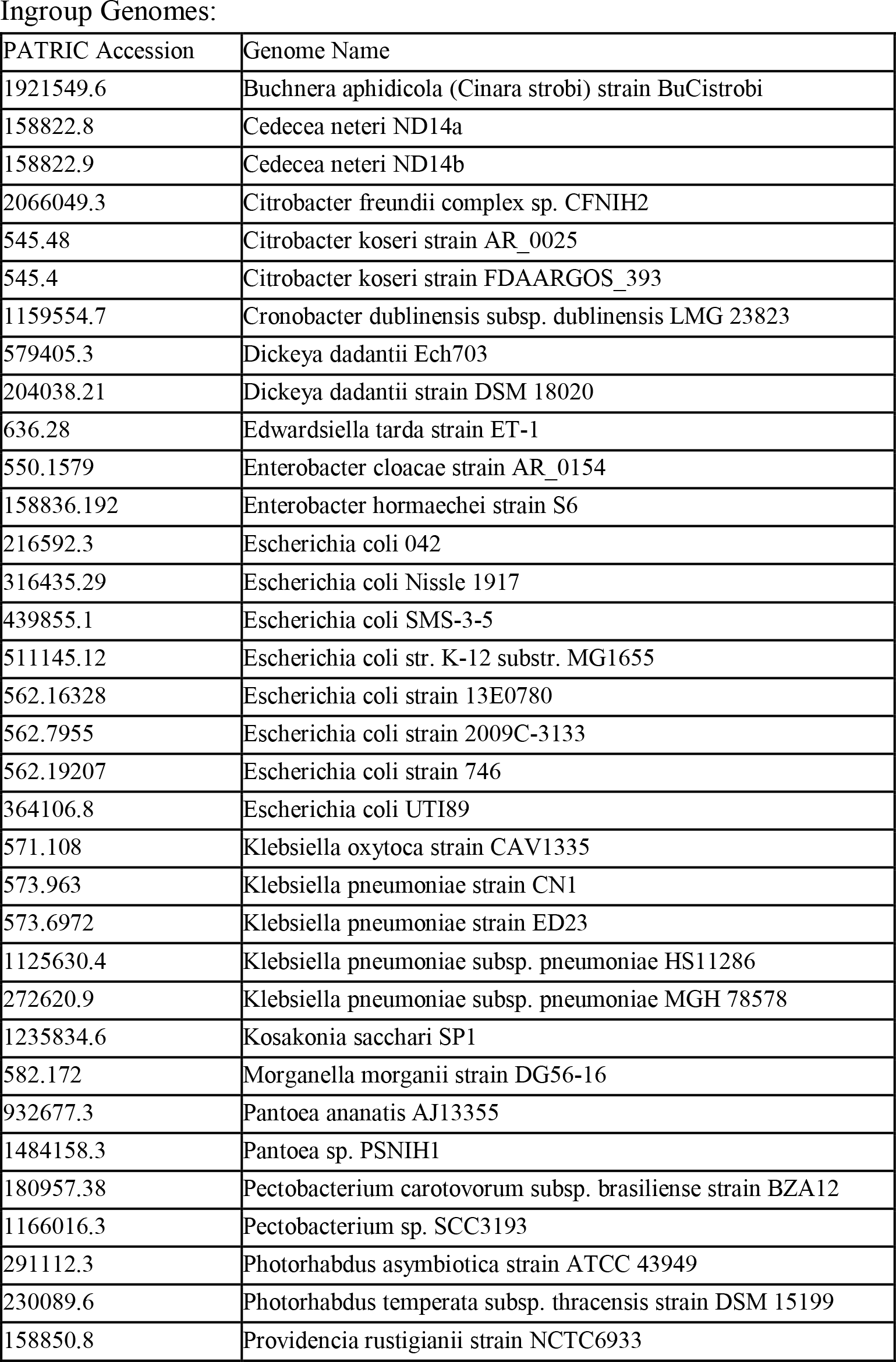
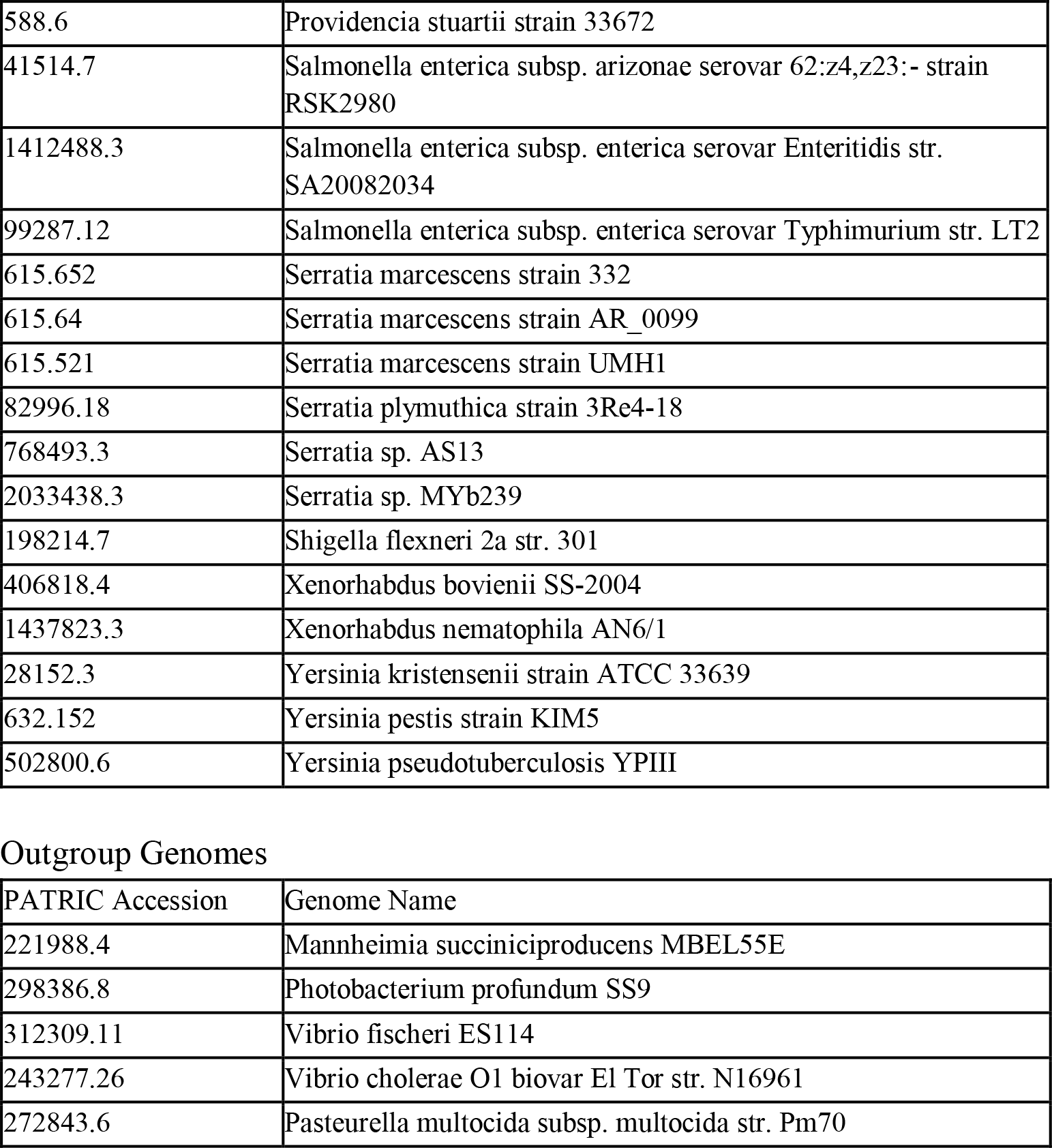
Input genomes for phylogenetic tree construction. List of 50 manually selected enterobacterial genomes from various genera with different distributions of BGC types used as in-group for construction of phylogenetic tree using maximum likelihood algorithm. List of five genomes from neighboring clades of Gammaproteobacteria used as out-group during phylogenetic tree construction.

**Table S3:**
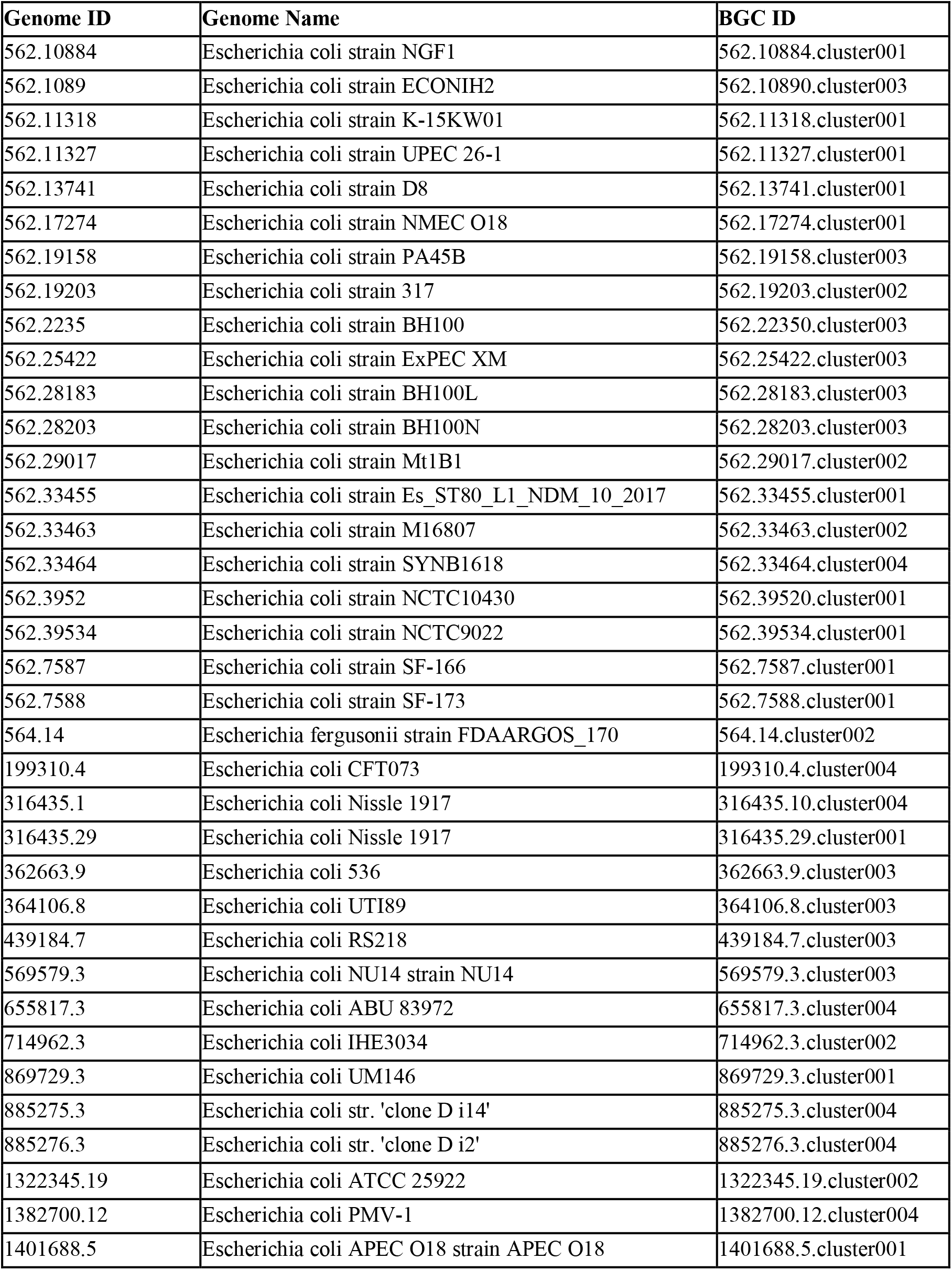
List of 36 *Escherichia* genomes with colibactin BGC that are selected for pangenome analysis.

| Genome ID | Genome Name | BGC ID |
| --- | --- | --- |
| 562.10884 | *Escherichia coli* strain NGF1 | 562.10884.cluster001 |
| 562.1089 | *Escherichia coli* strain ECONIH2 | 562.10890.cluster003 |
| 562.11318 | *Escherichia coli* strain K-15KW01 | 562.11318.cluster001 |
| 562.11327 | *Escherichia coli* strain UPEC 26-1 | 562.11327.cluster001 |
| 562.13741 | *Escherichia coli* strain D8 | 562.13741.cluster001 |
| 562.17274 | *Escherichia coli* strain NMEC O18 | 562.17274.cluster001 |
| 562.19158 | *Escherichia coli* strain PA45B | 562.19158.cluster003 |
| 562.19203 | *Escherichia coli* strain 317 | 562.19203.cluster002 |
| 562.2235 | *Escherichia coli* strain BH100 | 562.22350.cluster003 |
| 562.25422 | *Escherichia coli* strain ExPEC XM | 562.25422.cluster003 |
| 562.28183 | *Escherichia coli* strain BH100L | 562.28183.cluster003 |
| 562.28203 | *Escherichia coli* strain BH100N | 562.28203.cluster003 |
| 562.29017 | *Escherichia coli* strain Mt1B1 | 562.29017.cluster002 |
| 562.33455 | *Escherichia coli* strain Es_ST80_L1_NDM_10_2017 | 562.33455.cluster001 |
| 562.33463 | *Escherichia coli* strain M16807 | 562.33463.cluster002 |
| 562.33464 | *Escherichia coli* strain SYNB1618 | 562.33464.cluster004 |
| 562.3952 | *Escherichia coli* strain NCTC10430 | 562.39520.cluster001 |
| 562.39534 | *Escherichia coli* strain NCTC9022 | 562.39534.cluster001 |
| 562.7587 | *Escherichia coli* strain SF-166 | 562.7587.cluster001 |
| 562.7588 | *Escherichia coli* strain SF-173 | 562.7588.cluster001 |
| 564.14 | *Escherichia fergusonii* strain FDAARGOS_170 | 564.14.cluster002 |
| 199310.4 | *Escherichia coli* CFT073 | 199310.4.cluster004 |
| 316435.1 | *Escherichia coli* Nissle 1917 | 316435.10.cluster004 |
| 316435.29 | *Escherichia coli* Nissle 1917 | 316435.29.cluster001 |
| 362663.9 | *Escherichia coli* 536 | 362663.9.cluster003 |
| 364106.8 | *Escherichia coli* UTI89 | 364106.8.cluster003 |
| 439184.7 | *Escherichia coli* RS218 | 439184.7.cluster003 |
| 569579.3 | *Escherichia coli* NU14 strain NU14 | 569579.3.cluster003 |
| 655817.3 | *Escherichia coli* ABU 83972 | 655817.3.cluster004 |
| 714962.3 | *Escherichia coli* IHE3034 | 714962.3.cluster002 |
| 869729.3 | *Escherichia coli* UM146 | 869729.3.cluster001 |
| 885275.3 | *Escherichia coli* str. 'clone D i14' | 885275.3.cluster004 |
| 885276.3 | *Escherichia coli* str. 'clone D i2' | 885276.3.cluster004 |
| 1322345.19 | *Escherichia coli* ATCC 25922 | 1322345.19.cluster002 |
| 1382700.12 | *Escherichia coli* PMV-1 | 1382700.12.cluster004 |
| 1401688.5 | *Escherichia coli* APEC O18 strain APEC O18 | 1401688.5.cluster001 |

**Table S4:** Associated gene sets of colibactin gene BGC. List of 110 genes that are associatively present in Colibactin containing *Escherichia* genomes. Regions are defined if two or more neighbouring genes are present in associated gene sets. #clb represents the portion of colibactin possessing genomes with presence of a particular gene, which is 1 for all genes. #no_pks represent portion of genomes with corresponding gene, which is less than 0.1 here. Additionally, corresponding gene accession IDs for well-characterized strain *E. coli* CFT073 are listed. The background for important gene regions included in Fig. 4 is colored.

| Region | Gene ID (PATRIC Accession) | #clb | #ybt | #PKS distant | #no_pks | pan_gene_id | Product (PATRIC Annotation) | CFT073 gene ID |
| --- | --- | --- | --- | --- | --- | --- | --- | --- |
| 1 | fig\|1322345.19.peg.265\|DR76_253 | 1 | 0.41 | 0.12 | 0.03 | yfcQ | Minor fimbrial subunit StfF | c2880 |
| 1 | fig\|1322345.19.peg.266\|DR76_254 | 1 | 0.12 | 0.08 | 0.03 | yfcR | Minor fimbrial subunit StfE | c2881 |
| 1 | fig\|1322345.19.peg.267\|DR76_255 | 1 | 0.44 | 0.12 | 0.05 | yfcS | Periplasmic fimbrial chaperone StfD | c2882 |
| 1 | fig\|1322345.19.peg.268\|DR76_256 | 1 | 0.44 | 0.12 | 0.05 | yfcU | Fimbriae usher protein StfC | c2883 |
| 1 | fig\|1322345.19.peg.269\|DR76_257 | 1 | 0.44 | 0.12 | 0.04 | yfcV | Major fimbrial subunit StfA | c2884 |
| NA | fig\|1322345.19.peg.410\|DR76_398 | 1 | 0.57 | 0.29 | 0.07 | yfgJ | Uncharacterized protein YfgJ | c3032 |
| 2 | fig\|1322345.19.peg.516\|DR76_506 | 1 | 0.16 | 0.04 | 0.03 | group_12635 | hypothetical protein | c4897 |
| 2 | fig\|1322345.19.peg.519\|DR76_509 | 1 | 0.16 | 0.08 | 0.03 | tsx2 | Nucleoside-specific channel-forming protein Tsx precursor | c4894 |
| 3 | fig\|1322345.19.peg.636\|DR76_625 | 1 | 0.47 | 0.12 | 0.09 | group_5002 | hypothetical protein | c4770 |
| 3 | fig\|1322345.19.peg.637\|DR76_626 | 1 | 0.48 | 0.12 | 0.1 | rutB_2 | Isochorismatase (EC 3.3.2.1) | c4769 |
| 3 | fig\|1322345.19.peg.638\|DR76_627 | 1 | 0.48 | 0.12 | 0.1 | ybl188 | hypothetical protein | c4768 |
| 3 | fig\|1322345.19.peg.639\|DR76_628 | 1 | 0.48 | 0.12 | 0.1 | fdrA_3 | Membrane protein FdrA | c4767 |
| 3 | fig\|1322345.19.peg.640\|DR76_629 | 1 | 0.48 | 0.12 | 0.1 | ybl186 | Uncharacterized protein YahG | c4765 |
| 3 | fig\|1322345.19.peg.641\|DR76_630 | 1 | 0.48 | 0.12 | 0.1 | arcC | Carbamate kinase (EC 2.7.2.2) | c4764 |
| 3 | fig\|1322345.19.peg.642\|DR76_631 | 1 | 0.48 | 0.12 | 0.1 | atzF | Amidase | c4763 |
| 3 | fig\|1322345.19.peg.643\|DR76_632 | 1 | 0.48 | 0.12 | 0.1 | codB_2 | Cytosine/purine/uracil/thiamine/allantoin permease family protein | c4762 |
| 3 | fig\|1322345.19.peg.644\|DR76_633 | 1 | 0.48 | 0.12 | 0.1 | allS_6 | hypothetical protein | c4761 |
| 3 | fig\|1322345.19.peg.647\|DR76_636 | 1 | 0.43 | 0.12 | 0.02 | ptsG_2 | PTS system, IIC component / PTS system, IIB component | c4758 |
| 3 | fig\|1322345.19.peg.648\|DR76_637 | 1 | 0.43 | 0.12 | 0.02 | hxlB | related to 6-phospho-3-hexuloisomerase | c4757 |
| 3 | fig\|1322345.19.peg.651\|DR76_640 | 1 | 0.16 | 0.08 | 0.02 | group_13759 | hypothetical protein | c4754 |
| 4 | fig\|1322345.19.peg.808\|DR76_803 | 1 | 0.43 | 0.12 | 0.01 | group_1819 | hypothetical protein | c4581 |
| 4 | fig\|1322345.19.peg.809 | 1 | 0.15 | 0.08 | 0.01 | group_3328 | hypothetical protein | c4580 |
| 5 | fig\|1322345.19.peg.836\|DR76_826 | 1 | 0.53 | 0.17 | 0.06 | licR | Transcription antiterminator, BglG family | c4488 |
| 5 | fig\|1322345.19.peg.837\|DR76_827 | 1 | 0.53 | 0.17 | 0.06 | fruA_1 | PTS system, IIA component | c4487 |
| 5 | fig\|1322345.19.peg.838\|DR76_828 | 1 | 0.53 | 0.17 | 0.06 | manP | PTS system, IIB component | c4486 |
| 5 | fig\|1322345.19.peg.839\|DR76_829 | 1 | 0.53 | 0.17 | 0.06 | fruA_2 | PTS system, IIC component | c4485 |
| 5 | fig\|1322345.19.peg.840\|DR76_830 | 1 | 0.53 | 0.17 | 0.06 | kbaY | Fructose-bisphosphate aldolase class II (EC 4.1.2.13) homolog | c4484 |
| 5 | fig\|1322345.19.peg.841\|DR76_831 | 1 | 0.53 | 0.17 | 0.06 | fbaA_1 | Fructose-bisphosphate aldolase class II (EC 4.1.2.13) homolog | c4483 |
| 5 | fig\|1322345.19.peg.842\|DR76_832 | 1 | 0.53 | 0.17 | 0.06 | mak_2 | ROK family sugar kinase or transcriptional regulator | c4482 |
| 5 | fig\|1322345.19.peg.843\|DR76_833 | 1 | 0.53 | 0.17 | 0.06 | group_13463 | protein of unknown function DUF1498 | c4481 |
| 6 | fig\|1322345.19.peg.918\|DR76_905 | 1 | 0.39 | 0.17 | 0.05 | yiaM | 2,3-diketo-L-gulonate TRAP transporter small permease protein yiaM | c4399 |
| 6 | fig\|1322345.19.peg.919\|DR76_906 | 1 | 0.39 | 0.12 | 0.03 | group_13446 | Putative chemotaxis protein, resembles cheA | c4398 |
| NA | fig\|1322345.19.peg.1004 | 1 | 0.39 | 0.12 | 0.04 | group_5075 | hypothetical protein | c4311 |
| NA | fig\|1322345.19.peg.1092\|DR76_1070 | 1 | 0.18 | 0.12 | 0.04 | ycbS | Outer membrane usher protein fimD precursor | c4212 |
| 7 | fig\|1322345.19.peg.1266\|DR76_1244 | 1 | 0.55 | 0.17 | 0.09 | glpR_5 | Aga operon transcriptional repressor, DeoR family | c4020 |
| 7 | fig\|1322345.19.peg.1267\|DR76_1245 | 1 | 0.55 | 0.17 | 0.09 | gatY | Tagatose-1,6-bisphosphate aldolase GatY (EC 4.1.2.40) | c4018 |
| 7 | fig\|1322345.19.peg.1268\|DR76_1246 | 1 | 0.55 | 0.17 | 0.09 | rbsB_1 | ABC transporter, substrate-binding protein (cluster 2, ribose/xylose/arabinose/galactose) | c4017 |
| 7 | fig\|1322345.19.peg.1269\|DR76_1247 | 1 | 0.55 | 0.17 | 0.09 | rbsA_3 | Putative ribose/galactose/methyl galactoside import ATP-binding protein 1 (EC 3.6.3.17) | c4016 |
| 7 | fig\|1322345.19.peg.1270\|DR76_1248 | 1 | 0.55 | 0.17 | 0.09 | rbsC | ABC transporter, permease protein (cluster 2, ribose/xylose/arabinose/galactose) | c4015 |
| 7 | fig\|1322345.19.peg.1271\|DR76_1249 | 1 | 0.55 | 0.17 | 0.09 | iolC_2 | Putative sugar kinase, PfkB family protein (EC 2.7.1.-) | c4014 |
| 8 | fig\|1322345.19.peg.1476\|DR76_1449 | 1 | 0.51 | 0.25 | 0.09 | dsbB_1 | Inner membrane thiol:disulfide oxidoreductase, DsbB-like | c3787 |
| 8 | fig\|1322345.19.peg.1477\|DR76_1450 | 1 | 0.51 | 0.25 | 0.09 | dsbL | Periplasmic thiol:disulfide interchange protein, DsbA-like | c3786 |
| 8 | fig\|1322345.19.peg.1478\|DR76_1451 | 1 | 0.51 | 0.25 | 0.09 | asst | Putative arylsulfate sulfotransferase (EC 2.8.2.22) | c3785 |
| 9 | fig\|1322345.19.peg.1576\|DR76_1547 | 1 | 0.53 | 0.17 | 0.1 | kpsD | Capsular polysaccharide export system periplasmic protein KpsD | c3688 |
| 9 | fig\|1322345.19.peg.1578\|DR76_1549 | 1 | 0.53 | 0.17 | 0.1 | kpsF | D-arabinose-5-phosphate isomerase (EC 5.3.1.13) | c3686 |
| NA | fig\|1322345.19.peg.1579 | 1 | 0.47 | 0.12 | 0.09 | group_714 | hypothetical protein | c3685 |
| 10 | fig\|1322345.19.peg.1845\|DR76_1793 | 1 | 0.47 | 0.12 | 0.08 | group_13973 | Type VI secretion-related protein VasL | c3403 |
| 10 | fig\|1322345.19.peg.1846\|DR76_1794 | 1 | 0.43 | 0.12 | 0.07 | tssE | T6SS lysozyme-like component TssE | c3402 |
| 10 | fig\|1322345.19.peg.1847\|DR76_1795 | 1 | 0.43 | 0.12 | 0.07 | tssJ | T6SS secretion lipoprotein TssJ (VasD) | c3401 |
| 10 | fig\|1322345.19.peg.1848\|DR76_1796 | 1 | 0.43 | 0.12 | 0.07 | tssG | T6SS component TssF (ImpG/VasA) | c3400 |
| 10 | fig\|1322345.19.peg.1850\|DR76_1798 | 1 | 0.43 | 0.08 | 0.06 | tssA | T6SS component TssA (ImpA) | c3399 |
| 10 | fig\|1322345.19.peg.1860\|DR76_1808 | 1 | 0.47 | 0.12 | 0.08 | clpV | T6SS AAA+ chaperone ClpV (TssH) | c3392 |
| 10 | fig\|1322345.19.peg.1862\|DR76_1810 | 1 | 0.43 | 0.12 | 0.08 | yiaD_2 | Outer membrane protein | c3389 |
| 10 | fig\|1322345.19.peg.1863\|DR76_1811 | 1 | 0.43 | 0.12 | 0.08 | group_2482 | T6SS outer membrane component TssL (ImpK/VasF) | c3388 |
| 10 | fig\|1322345.19.peg.1864\|DR76_1812 | 1 | 0.43 | 0.12 | 0.08 | tssK | T6SS component TssK (ImpJ/VasE) | c3387 |
| 10 | fig\|1322345.19.peg.1865\|DR76_1813 | 1 | 0.47 | 0.12 | 0.09 | tssC | T6SS component TssC (ImpC/VipB) | c3386 |
| 10 | fig\|1322345.19.peg.1866\|DR76_1814 | 1 | 0.47 | 0.12 | 0.09 | tssB | T6SS component TssB (ImpB/VipA) | c3385 |
| 11 | fig\|1322345.19.peg.1938\|DR76_1884 | 1 | 0.4 | 0.12 | 0.01 | wrbA_2 | Multimeric flavodoxin WrbA | c3304 |
| 11 | fig\|1322345.19.peg.1939\|DR76_1885 | 1 | 0.4 | 0.12 | 0.01 | group_3562 | MBL-fold metallo-hydrolase superfamily | c3302 |
| NA | fig\|1322345.19.peg.2100\|DR76_2057 | 1 | 0.42 | 0.12 | 0.03 | group_1921 | hypothetical protein | c4978 |
| 12 | fig\|1322345.19.peg.2150\|DR76_2105 | 1 | 0.4 | 0.17 | 0.04 | sucA | 2-oxoglutarate dehydrogenase E1 component (EC 1.2.4.2) | c5032 |
| 12 | fig\|1322345.19.peg.2151\|DR76_2106 | 1 | 0.4 | 0.17 | 0.04 | sucB_2 | Dihydrolipoamide succinyltransferase component (E2) of 2-oxoglutarate dehydrogenase complex (EC 2.3.1.61) | c5034 |
| 12 | fig\|1322345.19.peg.2152\|DR76_2107 | 1 | 0.4 | 0.17 | 0.04 | lpdA | Dihydrolipoamide dehydrogenase of 2-oxoglutarate dehydrogenase (EC 1.8.1.4) | c5035 |
| 12 | fig\|1322345.19.peg.2154\|DR76_2109 | 1 | 0.4 | 0.17 | 0.04 | sucD_2 | Succinyl-CoA ligase [ADP-forming] alpha chain (EC 6.2.1.5) | c5037 |
| 12 | fig\|1322345.19.peg.2155\|DR76_2110 | 1 | 0.4 | 0.17 | 0.04 | citT_2 | Membrane protein | c5038 |
| 12 | fig\|1322345.19.peg.2157\|DR76_2111 | 1 | 0.4 | 0.17 | 0.04 | dctD | hypothetical protein | c5040 |
| 12 | fig\|1322345.19.peg.2158\|DR76_2113 | 1 | 0.4 | 0.17 | 0.04 | dctB | hypothetical protein | c5041 |
| NA | fig\|1322345.19.peg.2318\|DR76_2272 | 1 | 0.43 | 0.12 | 0.03 | group_1280 | HipB protein @ Antitoxin HigA | c5295 |
| 13 | fig\|1322345.19.peg.2367\|DR76_2318 | 1 | 0.22 | 0.04 | 0.01 | argR_2 | Arginine pathway regulatory protein ArgR, repressor of arg regulon | c5346 |
| 13 | fig\|1322345.19.peg.2369\|DR76_2320 | 1 | 0.26 | 0.29 | 0.03 | argF | Ornithine carbamoyltransferase (EC 2.1.3.3) | c5348 |
| 13 | fig\|1322345.19.peg.2371\|DR76_2322 | 1 | 0.26 | 0.29 | 0.03 | arcA | Arginine deiminase (EC 3.5.3.6) | c5350 |
| 13 | fig\|1322345.19.peg.2372 | 1 | 0.19 | 0.04 | 0.01 | group_1549 | hypothetical protein | c5351 |
| NA | fig\|1322345.19.peg.2377\|DR76_2327 | 1 | 0.27 | 0.33 | 0.06 | yjgN | Inner membrane protein YjgN | c5357 |
| 14 | fig\|1322345.19.peg.2593\|DR76_2529 | 1 | 0.44 | 0.17 | 0.08 | group_3721 | hypothetical protein | c0021 |
| 14 | fig\|1322345.19.peg.2595\|DR76_2531 | 1 | 0.44 | 0.17 | 0.08 | group_5392 | hypothetical protein | c0023 |
| NA | fig\|1322345.19.peg.2700\|DR76_2628 | 1 | 0.4 | 0.12 | 0.01 | imm_2 | hypothetical protein | c0137 |
| 15 | fig\|1322345.19.peg.2726\|DR76_2653 | 1 | 0.15 | 0.08 | 0.03 | yadK | Uncharacterized fimbrial-like protein YadK | c0167 |
| 15 | fig\|1322345.19.peg.2727\|DR76_2654 | 1 | 0.14 | 0.08 | 0.02 | yadL | Uncharacterized fimbrial-like protein YadL | c0168 |
| 16 | fig\|1322345.19.peg.2873\|DR76_2800 | 1 | 0.03 | 0.17 | 0.05 | yagA | Mobile element protein | c2473 |
| 16 | fig\|1322345.19.peg.2880\|DR76_2806 | 1 | 0 | 0.04 | 0 | paaH | 3-hydroxyacyl-CoA dehydrogenase (EC 1.1.1.35) | c2467 |
| 16 | fig\|1322345.19.peg.2881\|DR76_2807 | 1 | 0 | 0.04 | 0 | clbE | hypothetical protein | c2466 |
| 16 | fig\|1322345.19.peg.2882\|DR76_2808 | 1 | 0 | 0.04 | 0 | clbF | Acyl-CoA dehydrogenase | c2464 |
| 16 | fig\|1322345.19.peg.2883\|DR76_2809 | 1 | 0 | 0.04 | 0 | clbG | Malonyl CoA-acyl carrier protein transacylase (EC 2.3.1.39) in polyketide synthesis | c2463 |
| 16 | fig\|1322345.19.peg.2885\|DR76_2811 | 1 | 0 | 0.04 | 0 | clbI | Polyketide synthase modules and related proteins | c2460 |
| 16 | fig\|1322345.19.peg.2891\|DR76_2817 | 1 | 0.03 | 0.04 | 0.02 | clbO | Polyketide synthase modules and related proteins | c2453 |
| 16 | fig\|1322345.19.peg.2893\|DR76_2819 | 1 | 0.03 | 0.04 | 0.02 | clbQ | Thioesterase | c2451 |
| 16 | fig\|1322345.19.peg.2895\|DR76_2821 | 1 | 0.73 | 0.96 | 0.01 | intA_1 | Integrase | c2449 |
| 16 | fig\|1322345.19.peg.2908\|DR76_2835 | 1 | 1 | 1 | 0.01 | fyuA | iron aquisition outermembrane yersiniabactin receptor (FyuA,Psn,pesticin receptor) @ Outer membrane receptor for ferric siderophore | c2436 |
| 16 | fig\|1322345.19.peg.2909\|DR76_2836 | 1 | 1 | 1 | 0.01 | ybtE | iron aquisition 2,3-dihydroxybenzoate-AMP ligase (EC 2.7.7.58,Irp5) @ 2,3-dihydroxybenzoate-AMP ligase (EC 2.7.7.58) of siderophore biosynthesis | c2433 |
| 16 | fig\|1322345.19.peg.2910\|DR76_2837 | 1 | 1 | 1 | 0.01 | ybtT | Iron aquisition yersiniabactin synthesis enzyme YbtT @ Thioesterase in siderophore biosynthesis gene cluster | c2432 |
| 16 | fig\|1322345.19.peg.2911\|DR76_2838 | 1 | 1 | 1 | 0.01 | ybtU | Yersiniabactin synthetase, thiazolinyil reductase component Irp3 @ Thiazolinyil imide reductase in siderophore biosynthesis gene cluster | c2430 |
| 16 | fig\|1322345.19.peg.2914\|DR76_2841 | 1 | 1 | 1 | 0.01 | ybtA | iron aquisition regulator (YbtA,AraC-like,required for transcription of FyuA/psn,Irp2) | c2423 |
| 16 | fig\|1322345.19.peg.2915\|DR76_2842 | 1 | 1 | 1 | 0.01 | ybtP | Iron siderophore ABC transporter, permease/ATP-binding protein YbtP | c2422 |
| 16 | fig\|1322345.19.peg.2916\|DR76_2843 | 1 | 1 | 1 | 0.01 | ybtQ | Iron siderophore ABC transporter, permease/ATP-binding protein YbtQ | c2421 |
| 16 | fig\|1322345.19.peg.2917\|DR76_2844 | 1 | 1 | 1 | 0.01 | ybtX | Putative signal transducer | c2420 |
| NA | fig\|1322345.19.peg.3470 | 1 | 0.4 | 0.12 | 0.05 | group_11581 | hypothetical protein | c1915 |
| NA | fig\|1322345.19.peg.3502\|DR76_3396 | 1 | 0.14 | 0.12 | 0.02 | group_1605 | T6SS PAAR-repeat protein | c1882 |
| 17 | fig\|1322345.19.peg.3570\|DR76_3457 | 1 | 0.49 | 0.17 | 0.05 | yibH_1 | Membrane fusion component of MSF-type tripartite multidrug efflux system | c1811 |
| 17 | fig\|1322345.19.peg.3571\|DR76_3458 | 1 | 0.49 | 0.17 | 0.06 | ripA_2 | Transcriptional regulator, AraC family | c1810 |
| NA | fig\|1322345.19.peg.3572 | 1 | 0.43 | 0.12 | 0.04 | group_2010 | Aminobenzoyl-glutamate transport protein | c1809 |
| NA | fig\|1322345.19.peg.4428\|DR76_4281 | 1 | 0.43 | 0.12 | 0.01 | ycf3 | TPR domain protein | c0807 |
| 18 | fig\|1322345.19.peg.4471\|DR76_4319 | 1 | 0.39 | 0.08 | 0.01 | glcR_2 | D-threonate utilization transcriptional regulator, LysR family | c0765 |
| 18 | fig\|1322345.19.peg.4472\|DR76_4320 | 1 | 0.39 | 0.08 | 0.01 | pdxA | D-threonate 4-phosphate dehydrogenase [decarboxylating] | c0764 |
| 18 | fig\|1322345.19.peg.4473\|DR76_4321 | 1 | 0.39 | 0.08 | 0.01 | group_6138 | D-threonate kinase | c0763 |
| 18 | fig\|1322345.19.peg.4474\|DR76_4322 | 1 | 0.39 | 0.08 | 0.01 | gbsB | Alcohol dehydrogenase (EC 1.1.1.1) | c0762 |
| 18 | fig\|1322345.19.peg.4475\|DR76_4323 | 1 | 0.39 | 0.08 | 0.01 | dapA_2 | Dihydrodipicolinate synthase family protein APECO1_1389 | c0761 |
| 18 | fig\|1322345.19.peg.4478\|DR76_4326 | 1 | 0.39 | 0.08 | 0.01 | tabA_1 | Protein YjgK, linked to biofilm formation | c0758 |
| NA | fig\|1322345.19.peg.4589\|DR76_4439 | 1 | 0.39 | 0.17 | 0.03 | group_11857 | hypothetical protein | c0638 |
| NA | fig\|1322345.19.peg.4825 | 1 | 0.43 | 0.08 | 0.03 | group_2147 | hypothetical protein | c0398 |
| NA | fig\|1322345.19.peg.4852 | 1 | 0.18 | 0.12 | 0.02 | group_2151 | hypothetical protein | c0368 |

**Data S1: Input dataset of enterobacterial genomes from PATRIC database**

List of 2639 genomes downloaded with all the associated metadata that is present in the PATRIC database (sheet: downloaded_genomes). List of 12 genomes that are removed from the analysis after quality assessment (Methods)(sheet: removed_genomes). Overview of the N50 and N90 statistics (sheet: assembly_stats_N50). Final list of genomes in the curated set used for analysis (sheet: input_genomes). List of strains with more than 1 genome entries in the database (sheet: strains_with_multiple_genomes).

**Data S2: Secondary metabolite BGCs detected across 2627 genomes using antiSMASH**

List of 8604 BGCs detected using antiSMASH with details of BGC type, position, genome data and contig data where BGCs are found (sheet: detected_bgcs). List of 99 BGCs on the contig edge that are removed from the analysis (Methods) (sheet: removed_bgcs_on_contig_edge).

**Data S3: Sequence-based similarity network of BGCs**

List of 1784 BGC nodes from families 8 to 212 leading to the similarity network displayed in Fig. 2 (sheet: node_table_small_families). The various sequence similarity distances generated using BiG-SCAPE, the raw distance metric was used to define edges of the network (sheet: edge_table_small_famillies). List of BGCs in top 7 largest families (sheet: large_families_cluster_list).

**Data S4: Similarity network analysis of colibactin BGCs across genera**

List of 67 colibactin BGCs detected across genera and families detected using BiG-SCAPE (sheet: colibactin_bgcs). The various sequence similarity distances generated using BiG-SCAPE, the raw distance metric was used to define edges of the network (sheet: edge_table_colibactin_bgcs). Four of the BGCs from different genera and different families detected here are visualized in Fig. 3.

**Data S5: Pangenome analysis of 36 *Escherichia* genomes: Roary output data**

Total list of 14017 genes detected in the pangenome of 36 *Escherichia* genomes reconstructed using Roary software. Gene presence-absence tables across 36 genomes (sheet: gene_presence_absence/boolean). Clusters of homolog genes forming 14017 gene families in the pangenome (sheet: clustered_proteins).

**Data S6: Homology comparison of the core genome of colibactin containing Escherichia against all Escherichia genomes**

Percent identity score of bidirectional best blast hits of 2937 genes from the core genome of colibactin containing *Escherichia* against 782 genomes of *Escherichia*.

